# Robustness and applicability of functional genomics tools on scRNA-seq data

**DOI:** 10.1101/753319

**Authors:** Christian H. Holland, Jovan Tanevski, Jan Gleixner, Manu P. Kumar, Elisabetta Mereu, Javier Perales-Patón, Brian A. Joughin, Oliver Stegle, Douglas A. Lauffenburger, Holger Heyn, Bence Szalai, Julio Saez-Rodriguez

## Abstract

Many tools have been developed to extract functional and mechanistic insight from bulk transcriptome profiling data. With the advent of single-cell RNA sequencing (scRNA-seq), it is in principle possible to do such an analysis for single cells. However, scRNA-seq data has characteristics such as drop-out events, low library sizes and a comparatively large number of samples/cells. It is thus not clear if functional genomics tools established for bulk sequencing can be applied to scRNA-seq in a meaningful way. To address this question, we performed benchmark studies on *in silico* and *in vitro* single-cell RNA-seq data. We included the bulk-RNA tools PROGENy, GO enrichment and DoRothEA that estimate pathway and transcription factor (TF) activities, respectively, and compared them against the tools AUCell and metaVIPER, designed for scRNA-seq. For the *in silico* study we simulated single cells from TF/pathway perturbation bulk RNA-seq experiments. Our simulation strategy guarantees that the information of the original perturbation is preserved while resembling the characteristics of scRNA-seq data. We complemented the *in silico* data with *in vitro* scRNA-seq data upon CRISPR-mediated knock-out. Our benchmarks on both the simulated and real data revealed comparable performance to the original bulk data. Additionally, we showed that the TF and pathway activities preserve cell-type specific variability by analysing a mixture sample sequenced with 13 scRNA-seq different protocols. Our analyses suggest that bulk functional genomics tools can be applied to scRNA-seq data, outperforming dedicated single cell tools. Furthermore we provide a benchmark for further methods development by the community.

## Background

Gene expression profiles provide a blueprint of the status of cells. Thanks to diverse high-throughput techniques, such as microarrays and RNA-seq, expression profiles can be collected relatively easily, and are hence very common. To extract functional and mechanistic information from these profiles, many tools have been developed, that can, for example, estimate the status of molecular processes such as the activity of pathways or transcription factors (TFs). These functional genomics tools are broadly used and belong to the standard toolkit to analyze expression data [1–3].

Functional genomics tools typically combine prior knowledge with a statistical method to gain functional and mechanistic insights from omics data. In the case of transcriptomics, prior knowledge is typically rendered as gene sets containing genes belonging to, e.g., the same biological process or to the same Gene Ontology (GO) annotation. The Molecular Signature Database (MSigDB) is one of the largest collections of curated and annotated gene sets [4]. Statistical methods are as abundant as the different types of gene sets. Among them, the most commonly used are over-representation analysis (ORA) [5] and Gene Set Enrichment Analysis (GSEA) [6]. Still, there is a growing number of statistical methods spanning from simple linear models to advanced machine learning methods [7,8].

Recent technological advances in single-cell RNA-seq (scRNA-seq) enable the profiling of gene expression at the individual cell level [9]. Multiple methods have been developed, and they have experienced a dramatic improvement over recent years. However, they still have a number of limitations and biases, including low library size, and drop-outs. Bulk RNA-seq tools that focus on cell type identification and characterization as well as on inferring regulatory networks can be readily applied to scRNA-seq data [10]. This suggests that functional genomics tools should in principle be applicable to scRNA-seq data as well. However, it has not been investigated yet whether these limitations could distort and confound the results, rendering the tools not applicable to single-cell data.

In this paper, we benchmarked the robustness and applicability of different functional genomics methods on simulated and real scRNA-seq data. We focused on three tools for bulk and two for single cell RNA data. The bulk tools are PROGENy [11], DoRothEA [12] and classical GO enrichment analysis combining GO gene sets [13] with GSEA. PROGENy estimates the activity of 14 signaling pathways by combining corresponding gene sets with a linear model. DoRothEA is a collection of resources of TF’s targets (regulons) that can serve as gene sets for TF activity inference. For this study we coupled DoRothEA with the method VIPER [14] as it incorporates the mode of regulation of each TF-target interaction. Both PROGENy’s and DoRothEA’s gene sets are based on observing the transcriptomic consequences (the ‘footprint’) of the processes of interest rather than the genes composing the process as gene sets [15]. This approach has been shown to be more accurate and informative in inferring the process’s activity [11,16]. The tools specifically designed for application on scRNA-seq data that we considered are AUCell [17] and metaVIPER [18]. We coupled AUCell with gene sets from DoRothEA and PROGENy that we hereafter refer to as D-AUCell and P-AUCell. Using DoRothEA with both VIPER and AUCell on scRNA-seq for TF activity inference allowed us to compare the underlying statistical methods more objectively. metaVIPER is an extension of VIPER which is based on the same statistical method, but relies on multiple TF-regulon resources such as tissue specific regulatory networks.

We first benchmarked the tools on simulated single cell transcriptome profiles. We found that on this *in silico* data the gene sets from DoRothEA and PROGENy can functionally characterize simulated single cells. We observed that the performance of the different tools is dependent on the used statistical method and properties of the data, such as library size or number of cells. We then used real scRNA-seq data upon CRISPR-mediated knock-out/knock-down of TFs [19,20] to assess the performance of DoRothEA’s gene sets. The results of this benchmark further supported our finding that functional genomics tools can provide accurate mechanistic insights into single cells. We observed different performance by the different tool on this task dependent on the statistical approach used. Finally, we demonstrated the utility of the tools for pathway and TF activity estimation on recently published data profiling a complex sample with 13 different scRNA-seq technologies [21]. Here, we showed that summarizing gene expression into TF and pathway activities preserves cell type specific information. Collectively, our results suggest that the bulk based functional analysis tools DoRothEA and PROGENy outperform the single cell tools AUCell and metaVIPER. Although on scRNA-seq data DoRothEA and PROGENy were less accurate than on bulk RNA-seq, we were still able to extract relevant functional insights from scRNA-seq data.

## Results

### Robustness of bulk RNA based functional genomics tools against low gene coverage

Single-cell RNA-seq profiling is hampered by low gene coverage due to drop-out events [22]. In our first analysis we focused solely on the low gene coverage aspect and whether tools designed for bulk can deal with it. Specifically, We aimed to explore how DoRothEA, PROGENy and GO gene sets combined with GSEA (GO-GSEA) can handle low gene coverage in general, independently of other artefacts and characteristics from scRNA-seq protocols. Thus, we conducted this benchmark using bulk transcriptome benchmark data. In these studies, TFs and pathways are perturbed experimentally, and the transcriptome profile is measured before and after the perturbation. These experiments can be used to benchmark tools for TF/pathway activity estimation, as they should estimate correctly the change in the perturbed TF or pathway. The use of these datasets allowed us to systematically control for the gene coverage (see Methods). The workflow consisted in four steps (Fig. S1a). In the first step we summarized all perturbation experiments into a matrix of contrasts (with genes in rows and contrasts in columns) by differential gene expression analysis. Subsequently, we randomly replaced, independently for each contrast, logFC values with 0 so that we obtain a predefined number of “covered” genes with a logFC unequal to zero. Accordingly, a gene with a logFC = 0 was considered as missing/not covered. Afterwards we applied DoRothEA, PROGENy and GO-GSEA on the contrast matrix, subsetted only to those experiments which are suitable for the corresponding tool: TF perturbation for DoRothEA and pathway perturbation for PROGENy and GO-GSEA. We finally evaluate the global performance of the methods with Receiver operating characteristic (ROC) and precision recall (PR) curves (see Methods). This process was repeated 25 times to account for stochasticity effects during inserting zeros in the contrast matrix (see Methods).

DoRothEA’s TFs are accompanied by an empirical confidence level indicating the confidence in their regulons, ranging from A (most confident) to E (less confident) (see Methods). For this benchmark we included only TFs with confidence level A and B (denoted as DoRothEA (AB)) as this combination has a reasonable tradeoff between coverage and performance [12]. In general, the performance of DoRothEA dropped as gene coverage decreased. While it showed reasonable prediction power with all available genes (AUROC of 0.690), it approached almost the performance of a random model (AUROC of 0.5) when only 500 genes were covered (mean AUROC of 0.547; Fig. 1a, and similar trend with AUPRC, Fig. S1c).

**Fig. 1:**
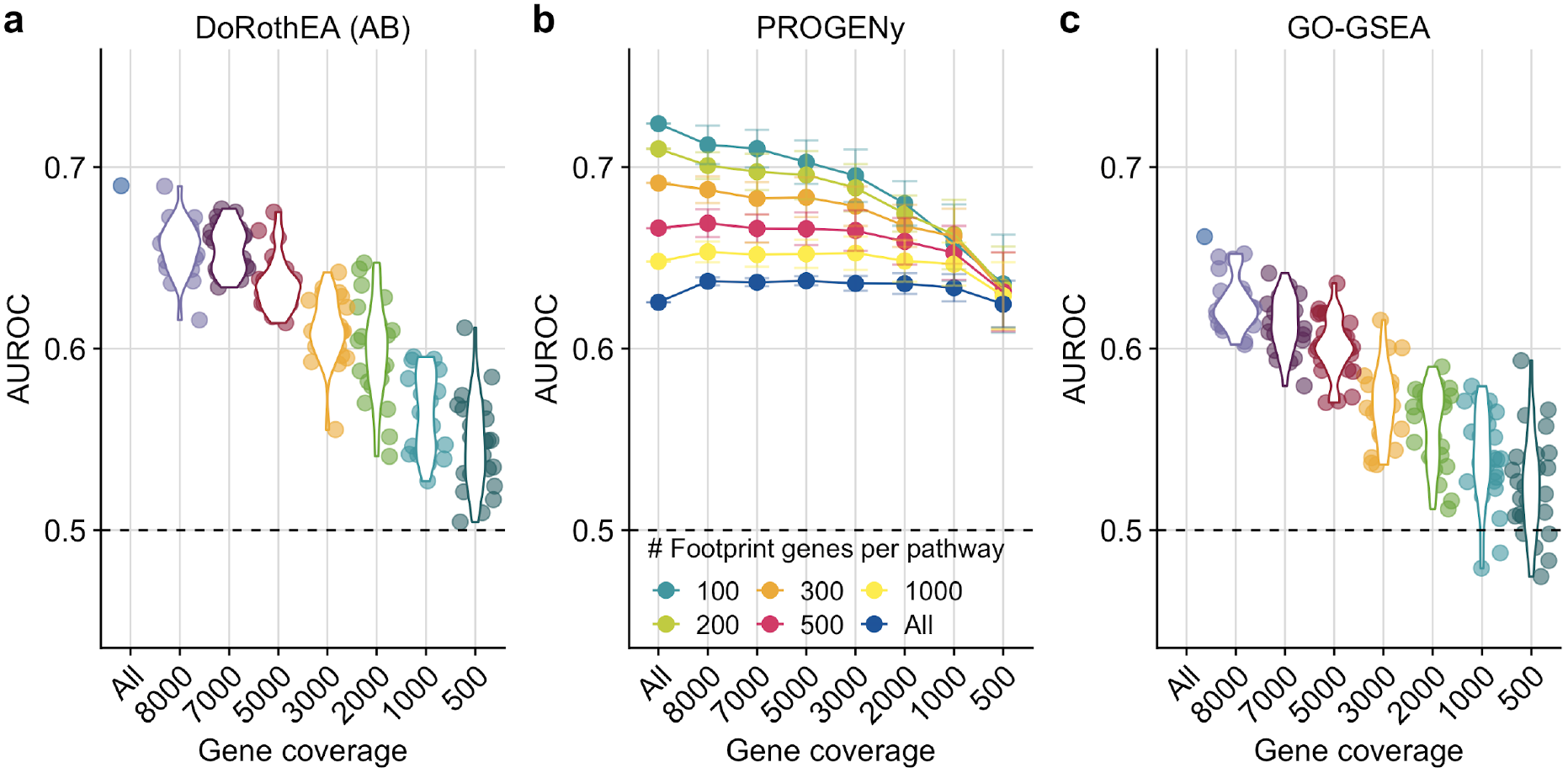
Testing the robustness of DoRothEA (AB), PROGENy and GO-GSEA against low gene coverage. **a** DoRothEA (AB) performance (Area under ROC curve, AUROC) versus gene coverage. **b** PROGENy performance (AUROC) for different number of footprint genes per pathway versus gene coverage. **c** Performance (AUROC) of GO-GSEA versus gene coverage. Dashed line indicates the performance of a random model.

We next benchmarked pathway activities estimated by PROGENy and GO-GSEA. In the original PROGENy framework, 100 footprint genes are used per pathway to compute pathway activities by default, as it has been shown that this leads to the best performance on bulk samples [11]. However, one can extend the footprint size to cover more genes of the expression profiles. We reasoned that this might counteract low gene coverage, and implemented accordingly different PROGENy versions (see Methods). With the default PROGENy version (100 footprint genes per pathway) we observed a clear drop in performance with decreasing gene coverage, even though less drastic than for DoRothEA (from AUROC of 0.724 to 0.636; Fig. 1b; similar trends with AUPRC; Fig. S1d). As expected, PROGENy performed the best with 100 footprint genes per pathway when there is complete gene coverage. The performance differences between the various PROGENy versions shrank with decreasing gene coverage. This suggests that increasing the number of footprint genes can help to counteract low gene coverage. To provide a fair comparison between PROGENy and GO-GSEA we used only those 14 GO terms that match the 14 PROGENy pathways (Fig. S1b). In general GO-GSEA showed weaker performance than PROGENy. The decrease in performance was more prominent as gene coverage decreased (from AUROC of 0.662 to 0.525; Fig. 1c and similar trend with AUPRC, Fig. S1e). With a gene coverage of less than 2000 genes, GO-GSEA performance was no better than random.

In summary, this first benchmark provided insight into the general robustness of the bulk based tools DoRothEA, PROGENy and GO-GSEA with respect to low gene coverage. DoRothEA performed reasonably well down to a gene coverage of 2000 genes. The performance of all different PROGENy versions were robust across the entire gene coverage range tested. GO-GSEA showed a worse performance than PROGENy, especially in the low gene coverage range. Since DoRothEA and PROGENy showed promising performance in low gene coverage ranges, we decided to explore them on scRNA-seq data. Due to its poor performance, we did not include GO-GSEA in the subsequent analyses.

### Benchmark of bulk and single-cell functional genomics tools on simulated scRNA-seq data

For the following analyses we expanded the set of tools by the methods AUCell [17] and metaVIPER [18]. Both methods were developed specifically for scRNA-seq analysis and thus allow the comparison of bulk vs. single-cell based tools on scRNA-seq data. AUCell is a statistical method that assesses whether gene sets are enriched in the top quantile of a ranked gene signature (see Methods). We combined AUCell with DoRothEA’s and PROGENy’s gene sets (referred to as D-AUCell and P-AUCell, respectively). metaVIPER is an extension of VIPER and requires multiple gene regulatory networks instead of a single network. In our study we coupled 27 tissue specific gene regulatory networks with metaVIPER, which provides a single TF consensus activity score estimated across all networks (see Methods). To benchmark all these methods on single cells, ideally we would have scRNA-seq datasets after perturbations of TFs and pathways. However, these datasets, especially for pathways, are currently very rare. To perform a comprehensive benchmark study, we developed a strategy to simulate samples of single cells using bulk RNA-seq samples from TF/pathway perturbation experiments.

A major cause of drop-outs in single cell experiments is the abundance of transcripts in the process of reverse-transcription of mRNA to cDNA [22]. Thus, our simulation strategy was based on the assumption that genes with low expression are more likely to result in drop-out events.

The simulation workflow started by transforming read counts of a single bulk RNA-seq sample to transcripts per million (TPM), normalizing for gene length and library size. Subsequently, for each gene, we assigned a sampling probability by dividing the individual TPM values with the sum of all TPM values. These probabilities are proportional to the likelihood for a given gene not to “drop-out” when simulating a single cell from the bulk sample. We determined the library size by sampling from a normal distribution with mean equal to the desired library size. For every single cell, we sampled with replacement genes from the gene probability vector up to the determined library size. The number of individual gene samples denote the new gene count in the single cell. The number of simulated single cells from a bulk sample is a parameter of the simulation (Fig. 2a, see Methods). This simple workflow guaranteed that the information of the original bulk perturbation is preserved and scRNA-seq characteristics, such as, drop-outs, low library size and high number of samples/cells are introduced.

**Fig. 2:**
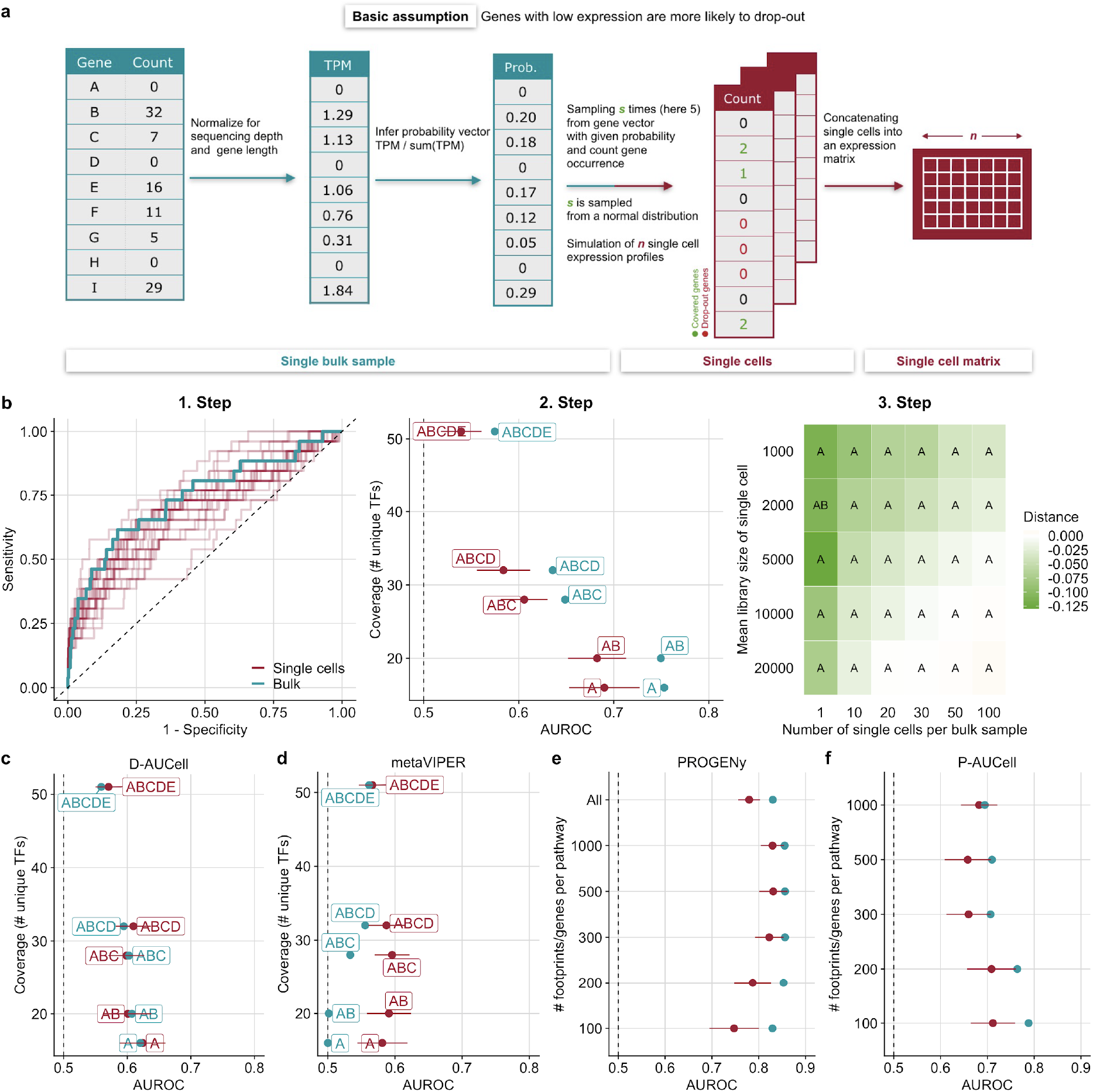
Benchmark result of various methods on simulated single cell data. **a** Simulation strategy of single cells from a RNA-seq bulk sample. **b** Example workflow of DoRothEA’s performance evaluation on simulated single cell. 1. step: ROC-curves of DoRothEA performance on simulated single cells (red lines) for 25 replicates of a specific parameter combination (number of cells = 10, mean library size = 5000) and on bulk data (blue line) including only TFs with confidence level A. Dashed line indicates the performance of a random model. 2. step: DoRothEA performance summarized as AUROC on simulated single cells (red lines) for a specific parameter combination (number of cells = 10, mean library size = 5000) and corresponding bulk data (blue line) vs TF coverage. Results are provided for different combinations of DoRothEA’s confidence levels (A,B,C,D,E). Error bars of AUROC values depict the standard deviation and correspond to different replicates of the given parameter combination. Dashed line indicates the performance of a random model. Step 3: Absolute difference across all confidence level combinations between AUROC of DoRothEA on single cells and AUROC of DoRothEA on bulk data for all parameter combinations. The letters within the tiles indicates which confidence level combination performs the best. The tile marked in red corresponds to the parameter setting used for previous plots (Step 1 and 2). **c** D-AUCell and **d** metaVIPER performance summarized as AUROC on simulated single cells (red lines) for a specific parameter combination (number of cells = 10, mean library size = 5000) and corresponding bulk data (blue line) vs TF coverage. Results are provided for different combinations of DoRothEA’s confidence levels (A,B,C,D,E). Error bars of AUROC values correspond to different replicates of the given parameter combination. Dashed line indicates the performance of a random model. **e and f** Performance result of **e** PROGENy and **f** P-AUCell on simulated single cells for a specific parameter combination (number of cells = 10, mean library size = 5000) and corresponding bulk in ROC space vs number of footprint genes per pathway. Error bars of AUROC values correspond to different replicates of the given parameter combination. Dashed line indicates the performance of a random model. **c d e f** Plots revealing the change in performance with varying simulation parameters (Step 3) are available in Supplementary Figure S6.

Our bulk RNA-seq samples comprised 97 single TF perturbation experiments targeting 52 TFs and 15 single pathway perturbation experiments targeting 7 pathways (Fig. S2a, S2b; see Methods). We repeated the simulation of numerous single cells from each bulk sample template to account for the stochasticity of the simulation procedure. We tested our simulation strategy by comparing the characteristics of the simulated cells to real single cells. We compared the count distribution (Fig. S3a and b), the relationship of mean and variance of gene expression (Fig. S3c and d) and the relationship of library size to number of detected genes (Fig. S3e and f). These comparisons suggested that our simulated single cells closely resemble real single cells and are thus suitable for benchmarking.

Unlike in our first benchmark, we applied the functional genomics tools directly on single samples/cells and built the contrasts at the level of pathway and TF activities (see Methods). We compared the performance of all tools to recover the perturbed TFs/pathways. We also considered the performance of the bulk based tools DoRothEA and PROGENy on the template bulk data as a baseline for comparison to their respective performance on the single cell data.

We show, as an example, the workflow of the performance evaluation for DoRothEA (Fig. 2b). As a first step we applied DoRothEA to single cells generated for one specific parameter combination (number of cells = 10, mean library size = 5000) and bulk samples, performed differential activity analysis (see Methods), and evaluated the performance with ROC and PR curves including only TFs with confidence level A. Each repetition of the simulation is depicted by an individual ROC curve, which shows the variance in performance of DoRothEA on simulated single cell data (Fig. 2b - 1. step). The variance decreases as the library size and the number of cells increase (which holds true for all tested tools; Fig. S4a-e). The shown ROC curves are summarized into a single AUROC value for bulk, and mean AUROC value for single cells. We performed this procedure also for different TF confidence level combinations and show the performance change in these values in relation to the TF coverage (Fig. 2b - 2. step). For both bulk and single cells, we observe a tradeoff between TF coverage and performance caused by including different TF confidence level combinations in the benchmark. This result is supported by both AUROC and AUPRC (Fig. S5a) and correspond to our previous findings [12]. The performance of DoRothEA on single cells does not reach the performance on bulk, though it can still recover TF perturbations on the simulated single cells reasonably well. This is especially true for the most confident TFs (AUROC of 0.690 for confidence level A and 0.682 for the confidence level combination AB). Finally we explore the effect of the library size and the number of cells on the performance by performing the previously described analysis for all combinations of library sizes and cell numbers. We computed the mean difference between AUROC scores of single cell and bulk data for all confidence level combinations. We observed a gradually decreasing differences when the size of the library and the number of cells increase (Fig. 2b - 3. step and Fig. S6a). Note, however, that the number of cells has a stronger impact on the performance than the mean library size. This analysis identified the best performing combination of DoRothEA’s TF confidence levels for different library sizes and number of single cells. Thus, the results can be used as recommendations for choosing the confidence levels on data from an experiment with comparable characteristics in terms of cell numbers and sequencing depths.

Similarly to DoRothEA, we also observed for D-AUCell a tradeoff between TF coverage and performance for both single cells and bulk samples when using the same parameter combination (Fig. 2c; similar trend with AUPRC Fig. S5b). The summarized performance across all confidence level combinations of D-AUCell on single cells slightly outperformed its performance on bulk samples (AUROC of 0.601 on single cells and 0.597 on bulk). This trend becomes more evident with increasing library size and number of cells (Fig. S6b).

For the benchmark of metaVIPER we assigned confidence levels to the tissue specific GTEx regulons based on DoRothEA’s gene set classification. This was done for consistency with DoRothEA and D-AUCell, even if there is no difference in confidence among them. Hence, for metaVIPER, we do not observe a tradeoff between TF coverage and performance (Fig. 2d; similar trend with AUPRC Fig. S5c). As opposed to D-AUCell, metaVIPER performed better on single cells than on bulk samples across all confidence level combinations (AUROC of 0.584 on single cells and 0.531 on bulk). This trend increased with increasing library size and number of cells (Fig. S6c). However, the overall performance of metaVIPER is worse than the performance of DoRothEA and D-AUCell. In summary, the bulk based tool DoRothEA performed the best on the simulated single cells followed by D-AUCell. metaVIPER performed slightly better that a random model.

For the benchmark of PROGENy we observed that it performed well across different number of footprint genes per pathway, with a peak at 500 footprint genes for both single cells and bulk (AUROC of 0.856 for bulk and 0.831 for single cells; Fig. 2e - similar trend with AUPRC Fig. S5d). A higher performance for single cell analysis with more than 100 footprint genes per pathway is in agreement with the previous general robustness study that suggested that a higher number of footprint genes can counteract low gene coverage. Increasing the library size and the number of cells improved the performance of PROGENy on single cells reaching almost the same performance as on bulk samples (Fig. S6d). For most parameter combinations, PROGENy with 500 or 1000 footprint genes per pathway yields the best performance.

For P-AUCell we observed a different pattern than for PROGENy as it worked best with 100 footprint genes per pathway for both single cells and bulk (AUROC of 0.788 for bulk and 0.712 for single cells; Fig. 2f - similar trends with AUPRC Fig. S5e). Similar to PROGENy, increasing the library size and the number of cells improved the performance, but not to the extent of its performance on bulk (Fig. S6e). For most parameter combinations P-AUCell with 100 or 200 footprint genes per pathway yielded the best performance.

In summary, both PROGENy and P-AUCell performed well on the simulated single cells, and PROGENy performed slightly better. For the pathway analysis the P-AUCell did not perform better on scRNA-seq than on bulk data. We then went on to perform a benchmark analysis on real scRNA-seq datasets.

### Benchmark of bulk and single-cell functional genomics tools on real scRNA-seq data

After showing that the gene sets from DoRothEA and PROGENy can handle low gene coverage and work reasonably well on simulated scRNA-seq data with different statistical approaches, we performed a benchmark on real scRNA-seq data. However, single cell transcriptome profiles of TF and pathway perturbations are very rare. To our knowledge there are no datasets of pathway perturbations on single cell level comprehensive enough for a robust benchmark of pathway analysis tools. For tools inferring TF activities the situation is better: recent studies combined CRISPR knock-outs/knock-down of TFs with scRNA-seq technologies [19,20], that can serve as potential benchmark.

The first dataset is based on the Perturb-seq technology, which contains 26 knock-out perturbations targeting 10 unique TFs after 7 and 13 days of perturbations (Fig. S7a) [19]. To explore the effect of perturbation time we divided the dataset into two sub datasets based on perturbation duration (Perturb-seq (7d) and Perturb-seq (13d)). The second dataset is based on CRISPRi protocol and contains 141 perturbation experiments targeting 50 unique TFs [20] (Fig. S7a). The datasets showed a variation in terms of drop-out rate, number of cells and sequencing depths (Fig. S7b).

To exclude bad or unsuccessful perturbations in case of CRISPRi experiments, we discarded experiments when the logFC of the targeted gene/TF was greater than 0 (12 out of 141; Fig. S7c). This quality control is important only in the case of CRISPRi, as it works on the transcriptional level. Perturb-seq (CRISPR knock-out) acts on the genomic level, so we can not expect a clear relationship between KO efficacy and transcript level of the target. Note that the logFC’s of both Perturb-seq sub datasets are in a narrower range in comparison to the logFCs of the CRISPRi dataset (Fig. S7d). The perturbation experiments which passed this quality check were used in the following analyses.

We evaluated the performance of DoRothEA, D-AUCell and metaVIPER on each benchmark dataset individually. We found that DoRothEA outperformed D-AUCell and metaVIPER across different combinations of DoRothEA confidence levels on Perturb-seq (7d) and CRISPRi dataset (Fig. 3a). metaVIPER did not perform better than a random model for these datasets. Interestingly, the performance of all three methods on the Perturb-seq (13d) dataset was very weak independently of DoRothEA’s confidence level (Fig. 3a). The captured trends are also reported in PR-space (Fig. S7e).

**Fig. 3:**
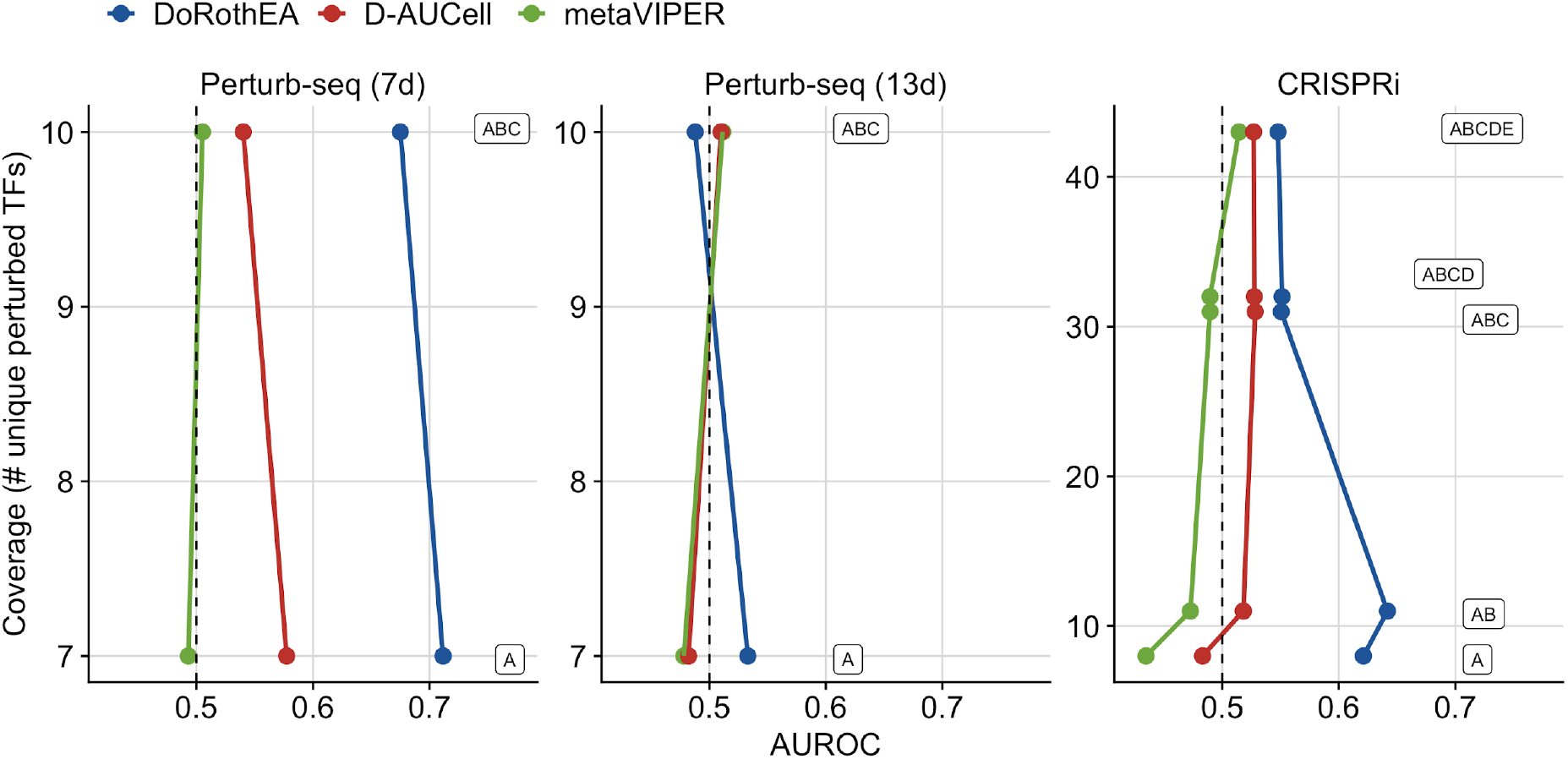
Benchmark result of VIPER on real single cell data. DoRothEA, D-AUCell and metaVIPER performance on all sub benchmark datasets in ROC space vs TF coverage split up by combinations of DoRothEA’s confidence levels (A-E).

In summary, these analyses suggested that DoRothEA is the best-performing method to recover TF perturbation at the single cell level on *in vitro* data.

### Application of bulk and single-cell functional genomics tools to samples of heterogeneous cell type populations (PBMC+HEK293T)

In our last analysis we wanted to test the performance of all tested tools in a more heterogeneous system that would illustrate a typical scRNA-seq data analysis scenario where multiple cell types are present. We used a dataset from the Human Cell Atlas project [23] that contains scRNA-seq profiles of human Peripheral blood mononuclear cells (PBMCs) and HEK293T with annotated cell types [21]. This dataset was analysed with 13 different scRNA-seq protocols (see Methods). In this study no ground truth (in contrast to the previous perturbation experiments) for TF and pathway activities were available. To evaluate the performance of all methods, we assessed the potential of TF and pathway activity estimations to cluster cells from the same cell type together based on *a priori* annotated cell types. We performed our analysis for each scRNA-seq technology separately to identify protocol-specific and protocol-independent trends. We assumed that the cell-type information should be preserved also on the reduced dimension space of TF / pathway activities, if these meaningfully capture the corresponding functional processes. Hence, we assessed how well the individual clusters correspond to the annotated cell types by a two-step approach. First we applied UMAP on different input matrices e.g. TF/pathway activities or gene expression and then we evaluated how well cells from the same cell type cluster together. We considered silhouette widths as a metric of cluster purity (see Methods). Silhouette widths derived from a set of highly variable genes (HVGs) set the baseline for the silhouette widths derived from pathway/TF activities. We identified the top 2000 HVGs with Seurat [24] using the selection method “vst” as it worked the best in our hands (Fig. S8). For both TF and pathway activity matrices the number of features available for dimensionality reduction using UMAP was substantially less (113 TFs and 14 pathways, respectively) than for a gene expression matrix containing the top 2000 HVGs. The number of available features for dimensionality reduction is different between HVGs, TFs, and pathways. To compare the cluster purity among these input features, we used positive and negative controls. The ‘positive control’ is a gene expression matrix with the top *n* HVGs and the negative control is a gene expression matrix with random *n* out of the 2000 HVGs (*n* equals 14 for pathway analysis and 113 for TF analysis).

Intuitively, each cell type should form a distinct cluster. However, some cell types are closely related, such as different T cells (CD4 and CD8) or Monocytes (CD14+ and FCGR3A+). Thus, we decided to evaluate the cluster purity at different levels of the cell-type hierarchy from fine-grained to coarse-grained. We start with the hierarchy level 0 where every cell type forms a distinct cluster and end with hierarchy level 4, where all PBMC cell types and all cell lines form a distinct cluster (Fig. 4a).

**Fig. 4:**
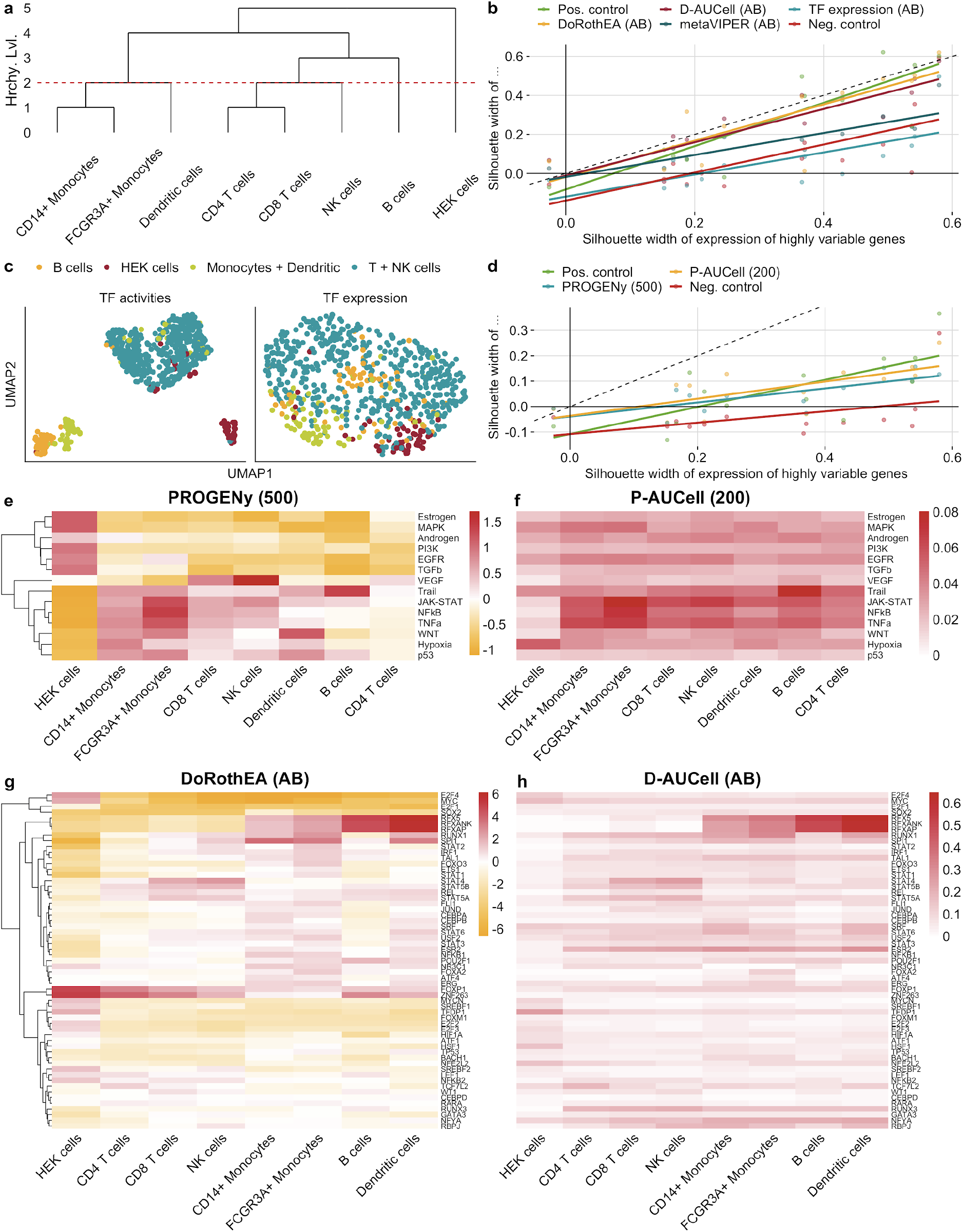
Application of DoRothEA/D-AUCell and PROGENy/P-AUCell) on a representative scRNA-seq dataset. **a** Dendrogram showing how cell lines/cell types are clustered together based on different hierarchy levels. Red dashed line marks the hierarchy level 2, where CD4 T cells, CD8 T cells and NK cells as well as CD14+ Monocytes, FCGR3A+ Monocytes and Dendritic cells are aggregated to a single cluster. **b,d** Comparison of cluster quality (clusters are defined by hierarchy level 2) between the top 2000 highly variable genes and **b** TF activity and TF expression, **d** pathway activities **c** UMAP plots of TF activities calculated with DoRothEA and corresponding TF expression measured by SMART-Seq2 protocol. **e and f** Pathway activities inferred from gene expression data (Quartz-Seq2) using **e** PROGENy and **f** (PROGENy AUCell). Pathway activities are summarized for each cell type/cell line separately. **g and h** Union of top 30 variable TF activities inferred from gene expression (Quartz-Seq2) using **g** DoRothEA and **h** D-AUCell summarized for each cell type/cell line separately.

To evaluate the performance of the TF activity inference methods and the utility of TF activity scores, we determined the cluster purity derived from TF activities (using only DoRothEA TFs with confidence level A and B), TF expression, positive and negative control. Both scRNA-seq protocols and matrices used for dimensionality reduction affected cluster purity significantly (2-way ANOVA p-values <2.2e-16 and 1.4e-10, respectively, p-values and estimations for corresponding linear model coefficients in Fig. S9a; see Methods). The cluster purity based on TF activities inferred using DoRothEA and D-AUCell did not differ significantly (Fig. 4b, corresponding plots for all hierarchy levels in Fig. S9b). In addition the cluster purity of both tools were not significantly worse that the purity based on all 2000 HVGs, though we observed a slight trend indicating a better cluster purity based on HVGs. This trend is expected due to the large difference of available features for dimensionality reduction. Instead a comparison to the positive and negative control is more appropriate. Both DoRothEA and D-AUCell performed comparably to the positive control but significantly better than the negative control across all scRNA-seq protocols (TukeyHSD post hoc test, adj. p-value of 1.05e-4 for DoRothEA and 5.7e-4 for D-AUCell). The cluster purity derived from metaVIPER was significantly worse than for DoRothEA (adj. p-value of 0.0423) and tend to be worse than D-AUCell (TukeyHSD post hoc test, adj. p-value of 0.130) as well. Also metaVIPER wasn’t better than than the negative control. Regardless of the underlying TF activity inference method, the cluster purity derived from TF activities outperformed the purity derived from TF expression (adj. p-value of 5.42e-6 for DoRothEA, 3.33-e5 for D-AUCell and 0.146 for metaVIPER). This underlines the advantage and relevance of using TF activities over the expression of the TF itself (Fig. 4c). With a comparable performance to a similar number of HVG and also to 2000 HVGs, we concluded that TF activities serve - independently of the underlying scRNA-seq protocol - as a complementary approach for cluster analysis that is based on generally more interpretable cell type marker.

To evaluate the performance of pathway inference methods and the utility of pathway activity scores we determined cluster purity with pathway matrices generated with different methods. We used 200 and 500 footprint genes per pathway for PROGENy and P-AUCell, respectively, since they provided the best performance in the previous analyses. As observed already before, both scRNA-seq protocols and matrices used for dimensionality reduction affected cluster purity significantly (2-way ANOVA p-values of 2.84e-7 and 1.13e-13, respectively, p values and estimations for corresponding linear model coefficients in Fig.S10b; see Methods). The cluster purity derived from pathway activity matrices is not significantly different PROGENy and P-AUCell, while worse than all HVGs (adj. p-value of 4.07e-10 for PROGENy and 4.59e-9 for P-AUCell; Fig. 4d, corresponding plots for all hierarchy levels in Fig. S9b). This is expected due to the large difference in the number of available features for dimensionality reduction (2000 HVGs vs 14 pathways). The cluster purity of both approaches was comparable to the positive control but significantly better than the negative control (adj. p-value of 0.077 for PROGENy and 0.013 for P-AUCell vs. negative control). In summary, this study indicated that the pathway activities contain relevant and cell-type specific information, even though they do not capture enough functional differences to be used for effective clustering analysis. Overall, the cluster purity of cells represented by the estimated pathway activities is worse than the cluster purity of cells represented by the estimated TF activities [21]. In addition we observed that input matrices derived from Quartz-Seq2 protocol yielded for hierarchy level 2 in significantly better cluster purity than all other protocols which is in agreement with the original study of of the PBMC + HEK293T data (Fig. S9a and S10a) [21].

TF and pathway activity scores are more interpretable than expression of single genes. Hence, we were interested to explore whether we could recover known cell-type specific TF and pathway activities from the PBMC data. We decided to focus on the dataset measured with Quartz-Seq2 as this protocol showed superior performance over all other platforms [21]. We calculated mean TF and pathway activity scores for each cell type using DoRothEA, D-AUCell, metaVIPER (all using only TFs with confidence levels A and B, Figure 4e, 4f and Supplementary Figure S11, respectively), PROGENy with 500 and P-AUCell with 200 footprint genes per pathway (Figure 4e-f). In agreement with the literature, we observed across both methods high activity of NFkB and TNFa in monocytes [25] and elevated Trail pathway activity in B cells (Fig. 4e-f) [26]. HEK cells, as expected from dividing cell lines, had higher activity of proliferative pathways (MAPK, EGFR and PI3K, Fig. 4e). These later pathway activity changes were only detected by PROGENy but not with AUCell, highlighting the importance of directionality information. Regarding TF activities, we observed high RFXAP, RFXANK and RFX5 activity (TFs responsible for MHCII expression) in monocytes, dendritic and B cells (the main antigen presenting cells of the investigated population [27]) (Fig. 4g-h). Myeloid lineage specific SPI1 activity [28] was observed in monocytes and dendritic cells. The high activity of repressor TF (where regulation directionality is important) FOXP1 in T lymphocytes [29] was only revealed by DoRothEA. Proliferative TFs like Myc and E2F4 had also high activity in HEK cells.

In summary, the analysis of this mixture sample demonstrated that summarizing gene expression into TF activities can preserve cell type specific information while drastically reducing the number of features. Hence, TF activity matrices could be considered as an alternative to full gene expression matrix for clustering analysis. We also showed that pathway activity matrices contain cell-type specific information, too, although we do not recommend using them for clustering analysis as the number of features is too low. In addition, we recovered known pathway/TF cell-type associations showing the importance of directionality and supporting the utility and power of the functional genomics tools DoRothEA and PROGENy.

## Discussion

In this paper we tested the robustness and applicability of functional genomics tools on scRNA-seq data. We included both bulk- and single-cell-based functional genomics tools that estimate either TF or pathway activities from gene expression data and for which well-defined benchmark data exist. The bulk based tools were DoRothEA, PROGENy and GO gene sets analysed with GSEA (GO-GSEA). The functional genomics tools specifically designed for the application in single cells were the statistical method AUCell combined with DoRothEA (D-AUCell) and PROGENy (P-AUCell) gene sets and metaVIPER.

We first explored the effect of low gene coverage in bulk data on the performance of the bulk based tools DoRothEA, PROGENy and GO-GSEA. We found that for all tools the performance dropped with decreasing gene coverage but at a different rate. While PROGENy was robust down to 500 covered genes, DoRothEA’s performance dropped markedly at 2000 covered genes. In addition, the results related to PROGENy suggested that increasing the number of footprint genes per pathway counteracted low gene coverage. GO-GSEA showed the strongest drop and did not perform better than a random guess below 2000 covered genes. Comparing the performance of both pathway analysis tools suggests that footprint based gene sets are superior over gene sets containing pathway members (e.g. GO gene sets) in recovering perturbed pathways. This observation is in agreement with previous studies conducted by us and others [11,30]. Given this fact and that GO-GSEA cannot handle low gene coverage (in our hands) we concluded that this approach is not suitable for scRNA-seq analysis. Hence, we decided to focus only on PROGENy as bulk based pathway analysis tool for the following analyses.

Afterwards, we benchmarked DoRothEA, PROGENy, D-AUCell, P-AUCell and metaVIPER on simulated single cells which we sampled from bulk pathway/TF perturbation samples. We showed that our simulated single cells possess characteristics comparable to real single cell data, supporting the relevance of this strategy. Different combinations of simulation parameters can be related to different scRNA-seq technologies. For each combination we provide a recommendation of how to use DoRothEA’s and PROGENy’s gene sets (in terms of confidence level combination or number of footprint genes per pathway) to yield the best performance. It should be noted that our simulation approach, as it is now, allows only the simulation of a homogenous cell population. This would correspond to a single cell experiment where the transcriptome of a cell line is profiled. In future work this simulation strategy could be adapted to account for a heterogeneous dataset which would resemble more realistic single cell datasets [31].

In terms of TF activity inference, DoRothEA performed best on the simulated single cells followed by D-AUCell and then metaVIPER. Both DoRothEA and D-AUCell shared DoRothEA’s gene set collection but applied different statistics. Thus, we concluded that, in our data, VIPER is more suitable to analyse scRNA-seq data than AUCell. The tool metaVIPER performed only slightly better than a random model and since it uses VIPER like DoRothEA the weak performance must be caused by the selection of the gene set resource. DoRothEA’s gene sets/TF regulons were constructed by integrating different types of evidence spanning from literature curated to predicted TF-target interactions. For metaVIPER we used 27 tissue specific gene regulatory networks constructed with ARACNe [32] thus containing only predicted TF-target interactions. The finding that especially the high confidence TFs regulons from DoRothEA outperform pure ARACNe regulons is in agreement with previous observations [12,33] and emphasizes the importance of combining literature curated resources with in silico predicted resources. Moreover, we hypothesize based on the pairwise comparison that for functional genomics analysis the choice of gene sets is of higher relevance than the choice of the underlying statistical method.

Related to pathway analysis, both PROGENy and P-AUCell performed well on the simulated single cells. The original framework of PROGENy uses a linear model that incorporates individual weights of the footprint genes, denoting the importance and also the sign of the contribution (positive/negative) to the pathway activity score. Those weights cannot be considered when applying AUCell with PROGENy gene sets. The slightly higher performance of PROGENy suggests that individual weights assigned to gene set members can improve the activity estimation of biological processes.

Especially in the benchmark of both TF analysis methods we observed that the D-AUCell and metaVIPER performed better on single cells than on the original bulk samples. This trend becomes more pronounced with increasing library size and number of cells. However, the bulk based tools perform better on single cells than the scRNA specific tools for both benchmarks.

Subsequently, we aimed to validate the aforementioned tools on real single cell data. While we could not find suitable benchmark data of pathway perturbations, we exploited two independent datasets of TF perturbations to benchmark the TF activity inference methods. These datasets combined CRISPR-mediated TF knock-out/knock-down (Perturb-Seq and CRISPRi) with scRNA-seq. It should be noted that pooled screenings of gene knock-outs with Perturb-seq suffer from an often faulty assignment of guide-RNA and single cell [34]. Those mislabeled data confound the benchmark as the groundtruth is not reliable. Nevertheless, we showed that DoRothEA’s gene sets were globally effective in inferring TF activity from single cell data with varying performance dependent on the used statistical method. As already shown in the *in silico* benchmark D-AUCell showed a weaker performance than DoRothEA, supporting that VIPER performs better than AUCell. Interestingly, metaVIPER’s performance was no better than random across all datasets. metaVIPER used the same statistical method as DoRothEA but different gene set resources. This further supports our hypothesis that the selection of gene sets is more important than the statistical method for functional genomics analysis.

Furthermore, the perturbation time had a profound effect on the performance of the tools: while DoRothEA and D-AUCell worked well for a perturbation duration of 6 (CRISPRi) and 7 days (Perturb-Seq (7d)), the performance dropped markedly for 13 days.

We reasoned that, within 13 days of perturbation, compensation effects are taking place at the molecular level that confound the prediction of TF activities. In addition, it is possible that cells without a gene edit outgrow cells with a successful knock-out after 13 days as the knock-out typically yield in a lower fitness and thus proliferation rate.

In summary, DoRothEA subsetted to confidence levels A and B performed the best on real scRNA-seq data but at the cost of the TF coverage. The results of the *in silico* and *in vitro* benchmark are in agreement. Accordingly, we believe that it is reasonable to assume that also PROGENy works on real data given the positive benchmark results on simulated data.

Finally, we applied our tools of interest to a mixture sample of PBMCs and HEK cells profiled with 13 different scRNA-seq protocols. We investigated to which extent pathway and TF matrices retain cell-type specific information, by evaluating how well cells belonging to the same cell type or cell type family cluster together in reduced dimensionality space. Given the lower numbers of features available for dimensionality reduction using TF and pathway activities, cell types could be recovered equally well as when using the same number of the top highly variable genes. In addition, we showed that cell types could be recovered more precisely using TF activities than TF expression, which is in agreement with previous studies [18]. This suggests that summarising gene expression as TF and pathway activities can lead to noise filtering, particularly relevant for scRNA-seq data. Though, TF activities performed better than pathway activities which is again attributed to the even lower number of pathways.

Our analysis suggested at different points that the performance of functional genomics tools is more sensitive to the selection of gene sets than the statistical methods. This hypothesis could be tested in future by decoupling functional genomics tools into gene sets and statistics. Benchmarking all possible combinations of gene sets and statistic (i.e. DoRothEA gene sets with a linear model or PROGENy gene sets with VIPER) would shed light on this question which we believe if of high relevance for the community.

## Conclusions

Our systematic and comprehensive benchmark study suggests that DoRothEA and PROGENy are effective in inferring TF and pathway activity from scRNA-seq data, outperforming tools specifically designed for scRNA-seq analysis. We showed the limits of both tools with respect to low gene coverage and also provided as part of the *in silico* benchmark recommendations on how to use DoRothEA’s and PROGENy’s gene sets in the best way dependent on the number of cells and mean library size. These two parameters are technology specific, so that our recommendations are transferable to various scRNA-seq protocols. Furthermore, we showed that TF and pathway activities are rich in cell type specific information with reduced amount of noise and provide an intuitive way of interpretation and hypothesis generation. We provide our benchmarks and code to the community for further assessment of methods for functional analysis.

## Methods

### PROGENy

PROGENy is a functional genomics tool which infers pathway activity for 14 signaling pathways (Androgen, Estrogen, EGFR, Hypoxia, JAK-STAT, MAPK, NFkB, PI3K, p53, TGFb, TNFa, Trail, VEGF and WNT) from gene expression data [11,35]. Pathway activity inference is based on gene sets comprising the top 100 most responsive genes upon corresponding pathway perturbation, which we refer to as footprint genes of a pathway.

### DoRothEA

DoRothEA is a data resource containing signed transcription factor (TF) - target interactions [12]. Those interactions were curated and collected from different types of evidence such as literature curated resources, ChIP-seq peaks, TF binding site motifs and interactions inferred directly from gene expression. Based on the number of supporting evidences each interaction is accompanied with an interaction confidence score ranging from A-E, with A being the most confidence interactions. In addition a summary TF confidence score is assigned (also from A-E) which is derived by and subsetted to the leading confidence level of its interactions. DoRothEA contains in total 470,711 interactions covering 1,396 TF targeting 20,238 unique genes. We use VIPER in combination with DoRothEA to estimate TF activities from gene expression data.

### VIPER

VIPER is a statistical framework which was developed to estimate protein activity from gene expression data using enriched regulon analysis performed by the algorithm aREA [14]. It requires information about (if possibly) signed interactions between a protein and its functional targets. In the original workflow this regulatory network was inferred from gene expression by the algorithm ARACNe [32]. However, it can be replaced by any other data resource reporting protein target interactions.

### Simulation of single cells

Let C be a vector representing counts per gene for a single bulk sample. C is normalized for gene length and library size resulting in vector B containing TPM values per gene. We assume that samples are obtained from homogenous cell populations and that the probability of a dropout event is proportional to the relative TPM of each measured gene in the bulk sample. Therefore, we define a discrete cumulative distribution function from the vector of gene frequencies 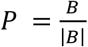. To simulate a single cell from this distribution, we draw and aggregate L samples by inverse transform sampling. L corresponds to the library size for the count vector of the simulated single cell. We draw L from a normal distribution 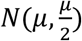.

To benchmark the robustness of the methods, we vary the number of cells sampled from a single bulk sample (1, 10, 20, 30, 50, 100) and the value of μ (1000, 2000, 5000, 10.000, 20.000). To account for stochasticity effects during sampling we repeat this analysis 25 times for each parameter combination.

Prior to normalization we discarded cells with a library size lower than 100. We normalized the count matrices of the simulated cells by using the R package *scran* (version 1.11.27) [36]. Contrast matrices were constructed by comparing cells originating from one of the perturbation bulk samples vs cells originating from one of the control bulk samples.

### Induction of artificial low gene coverage in bulk microarray data

We induce the reduction of gene coverage with inserting zeros on the contrast level. In detail we insert for each contrast separately randomly zeros until we obtained a predefined number of genes with a logFC unequal zero which we consider as “covered”/”measured” genes. We perform this analysis for a gene coverage of 500, 1000, 2000, 3000, 5000, 7000, 8000 and as reference all available genes. To account for stochasticity effects during inserting randomly zero we repeat this analysis 10 times for each gene coverage value.

### Application of PROGENy on single samples and contrasts

We applied PROGENy on matrices of single samples (genes in rows and either bulk samples or single cells in columns) containing normalized gene expression scores or on contrast matrices (genes in rows and summarized perturbation experiments into contrasts in columns) containing log fold changes. In case of single sample analysis the contrasts were built based on pathway activity matrices yielding the change in pathway activity (perturbed samples - control sample) summarized as logFC. Independent of input matrix we scaled each pathway to have a mean activity of 0 and a standard deviation of 1.

We build different PROGENy versions by varying the number of footprint genes per pathway (100, 200, 300, 500, 1000 or all which corresponds to ~29,000 genes).

### Application of VIPER on single samples and contrasts

We applied VIPER with DoRothEA as regulatory network resource on matrices of single samples (genes in rows and either bulk samples or single cells in columns) containing normalized gene expression scores scaled gene-wise to a mean value of 0 and standard deviation of 1 or on contrast matrices (genes in rows and summarized perturbation experiments into contrasts in columns) containing log fold changes. In case of single sample analysis the contrasts were built based on TF activity matrices yielding the change in TF activity (perturbed samples - control sample) summarized as logFC. TFs with less than 4 targets listed in the corresponding input matrix were discarded from the analysis. VIPER provides a NES enrichment score for each TF which we consider as a metric for the activity. We used the R package *viper* (version 1.17.0) [14] to run VIPER in combination with DoRothEA.

### Application of GSEA with GO gene sets on contrasts

We applied GSEA with gene sets on contrast matrices (genes in rows and summarized perturbation experiments into contrasts in columns) containing log fold changes that serve also a gene level statistic. We selected only those GO terms which map to PROGENy pathways in order to guarantee a fair comparison between both methods. For the enrichment analysis we used the R package *fgsea* (version 1.10.0) [37] with 1000 permutations per gene signature.

### Application of metaVIPER on single samples

We ran metaVIPER with 27 tissue specific gene regulatory networks which we constructed before for one of our previous studies [12]. Those tissue specific gene regulatory networks were derived using ARACNe [32] taking the database GTEx [38] as tissue specific gene expression sample resource. We applied metaVIPER on matrices of single samples (genes in rows and single cells in columns) containing normalized gene expression scores scaled gene-wise to a mean value of 0 and standard deviation of 1. Contrasts were built based on TF activity matrices yielding the change in TF activity (perturbed samples - control sample) summarized as logFC. TFs with less than 4 targets listed in the corresponding input matrix were discarded from the analysis. metaVIPER provides a NES integrated across all regulatory networks for each TF which we consider as a metric for the activity. We used the R package *viper* (version 1.17.0) [14] to run metaVIPER

### Application of AUCell with either DoRothEA or PROGENy gene sets on single samples

AUCell is a statistical method to determine specifically for single cells whether a given gene set is enriched at the top quantile of a ranked gene signature. Therefore AUCell determines the area under the recovery curve to compute the enrichment score. We defined the top quantile as the top 5 % of the ranked gene signature. We applied this method coupled with PROGENy and DoRothEA gene sets. Before applying this method with PROGENy gene sets we subsetted the footprint gene sets to contain only genes available in the provided gene signature. This guarantees a fair comparison as for the original PROGENy framework with a linear model the intersection of footprint (gene set) members and signature genes are considered. We applied AUCell with PROGENy and DoRothEA gene sets on matrices of single samples (genes in rows and single cells in columns) containing raw gene counts. Contrasts were built based on respective TF/pathway activity matrices yielding the change in TF/pathway activity (perturbed samples - control sample) summarized as logFC. For the AUCell analysis we used the R package *AUCell* (version 1.5.5) [17]

### Benchmarking process with ROC and PR metrics

To transform the benchmark into a binary setup, all activity scores of experiments with negative perturbation effect (inhibition/knockdown) are multiplied by −1. This guarantees, that TFs/pathways belong to a binary class either deregulated or not regulated and that the perturbed pathway/TF has in the ideal case the highest activity.

We performed the ROC and PR analysis with the R package *yardstick* (version 0.0.3; https://github.com/tidymodels/yardstick). For the construction of ROC and PR curves we calculated for each perturbation experiment pathway (or TF) activities using PROGENy (or VIPER). As each perturbation experiment targets either a single pathway (or TF) only the activity score of the perturbed pathway (or TF) is associated with the positive class (e.g. EGFR pathway activity score in an experiment where EGFR was perturbed). Accordingly the activity scores of all non-perturbed pathways (or TFs) belong to the negative class (e.g. EGFR pathway activity score in an experiment where JAK-STAT pathway was perturbed). Using these positive and negative classes Sensitivity / (1-Specificity) or Precision / Recall values were calculated at different thresholds of activity, producing the ROC / PRC curves.

### Collecting, curating and processing of microarray data

We extracted single pathway and single TF perturbation data profiled with classical microarrays from a previous study conducted by us [35]. We followed the same procedure of collection, curating and processing the data as described in the previous study.

### Collecting, curating and processing of bulk RNA-seq data

For the simulation of single cells we collected, curated and processed single TF and single pathway perturbation data profiled with bulk RNA-seq. We downloaded meta data of single TF perturbation experiments from the ChEA3 web-server (https://amp.pharm.mssm.edu/chea3/) [33]. Meta data of single pathway perturbation experiments were manually extracted by us from Gene Expression Omnibus (GEO) [39]. Count matrices for all those experiments were downloaded from ARCHS^4^ (https://amp.pharm.mssm.edu/archs4/) [40].

We normalized count matrices by first calculating normalization factors and second transforming count data to log2 counts per million (CPM) using the R packages *edgeR* (version 3.25.8) [41] and *limma* (version 3.39.18) [42], respectively.

### Collecting, curating and processing of scRNA-seq data for benchmark

To benchmark VIPER on real single cell data, we inspected related literature and identified two publications which systematically measure effects of transcription factors on gene expression in single cells:

Dixit et al. introduced Perturb-seq and measured the knockout-effects of 10 transcription factors on K562 cells 7 and 13 days after transduction [19]. We downloaded the expression data from GEO (GSM2396858 and GSM2396859) and sgRNA-cell mappings made available by author upon request in the files promoters_concat_all.csv (for GSM2396858) and pt2_concat_all.csv (for GSM2396859) on github.com/asncd/MIMOSCA. We did not consider the High MOI dataset due to the expected high number of duplicate sgRNA assignments. Cells were quality filtered based on expression, keeping the upper half of cells for each dataset. Only sgRNAs detected in at least 30 cells were used. For the day 7 dataset, 16507, and for day 13 dataset, 9634 cells remained for benchmarking.

Ryan at al. measured knockdown effects of 50 transcription factors implicated in human definitive endoderm differentiation using a CRISPRi variant of CROPseq in human embryonic stem cells 6 days after transduction [20]. We obtained data of both replicates from GEO (GSM3630200, GSM3630201), which include sgRNA counts next to the rest of the transcription. We refrained from using the targeted sequencing of the sgRNA in GSM3630202, GSM3630203 as it contained less clear mappings due to amplification noise. Expression data lacked information on mitochondrial genes and therefore no further quality filtering of cells was performed. From this dataset, only sgRNAs detected in at least 100 cells were used. A combined 5282 cells remained for benchmarking.

Analysis was limited to the 10000 most expressed genes for all three datasets. We normalized the count matrices for each individual dataset (Perturb-Seq (7d), Perturb-Seq (13d) and CRISPRi) separately by using the R package *scran* (version 1.11.27) [36].

### Collecting, curating and processing of scRNA-seq data from cell atlas project

This scRNA-seq dataset originates from a benchmark study of the Human Cell Atlas project [21]. At the time of writing this dataset is not publicly available but will be accessible from Gene Expression Omnibus in the near future (GSE133549). The dataset consists of a PBMC’s and a HEK293T sample which was analyzed with 13 different scRNA-seq technologies (CEL-Seq2, MARS-Seq, Quartz-Seq2, gmcSCRB-Seq, ddSEQ, ICELL8, C1HT-Small, C1HT-Medium, Chromium, Chromium(sn), Drop-seq, inDrop). Most cells are annotated with a specific cell type/cell line (CD4 T cells, CD8 T cells, NK cells, B cells, CD14+ Monocytes, FCGR3A+ Monocytes, Dendritic cells, Megakaryocytes, HEK cells). Cells without annotation were discarded for this analysis.

We normalized the count matrices for each technology separately by using the R package *scran* (version 1.11.27) [36].

### Dimensionality reduction with UMAP and assessment of cluster quality

We used the R package *umap* (version 0.2.0.0) calling the Python implementation of Uniform Manifold Approximation and Projection (UMAP) with the argument “method = ‘umap-learn’” to perform dimensionality reduction on various input input matrices (gene expression matrix, pathway/TF activity matrix, etc.). We assume that the dimensionality reduction will result in clustering of cells that corresponds well to the cell type/cell type family. To assess the validity of this assumption, we assigned a cell-type/cell family specific cluster id to each point in the low-dimensional space. We then defined a global cluster purity measure based on silhouette widths [43], which is a well known clustering quality measure.

Given the cluster assignments, in the low-dimensional space, for each cell the average distance (*a*) to the cells that belong to the same cluster is calculated. Then the smallest average distance (*b*) to all cells belonging to the newest foreign cluster is calculated. The difference, between the latter and the former indicates the width of the silhouette for that cell, i.e. how well the cell is embedded in the assigned cluster. To make the silhouette widths comparable, they are normalized by dividing the difference with the larger of the two average distances 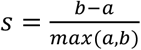. Therefore, the possible values for the silhouette widths lie in the range −1 to 1, where higher values indicate good cluster assignment, while lower values close to 0 indicate poor cluster assignment. Finally, the average silhouette width for every cluster is calculated, and averages are aggregated to obtain a measure of the global purity of clusters. For the silhouette analysis we used the R package *cluster* (version 2.0.8).

For statistical analysis of cluster quality, we fitted a linear model *score=f(scRNA-seq protocol + input matrix)*, where *score* corresponds to average silhouette width for a given scRNA-seq *protocol* - *input matrix* pair. *Protocol* and *input matrix* are factors, with reference level Quartz-Seq2 and positive control, respectively. We fitted two separate linear model for transcription factor and pathway activity inference methods. We report the estimates and p values for the different coefficients of these linear models. Based on these linear models, we performed a 2-way ANOVA, and pairwise comparisons using Tukey HSD post hoc test.

## Declarations

### Ethics approval and consent to participate

Not applicable

## Consent for publication

Not applicable

## Availability of data and material

The code to perform all presented studies is written in R [44] and is freely available on GitHub : https://github.com/saezlab/FootprintMethods_on_scRNAseq.

## Acknowledgments

We thank Aurélien Dugourd and Ricardo Ramirez-Flores for helpful discussions. We also thank Minoo Ashtiani for supporting the collection of single pathway perturbation experiments on bulk level.

## Funding

CHH is supported by the German Federal Ministry of Education and Research (BMBF)-funded project Systems Medicine of the Liver (LiSyM, FKZ: 031L0049). MPK, BAJ, DAL are supported by NIH Grant U54-CA217377. BS is supported by the Premium Postdoctoral Fellowship Program of the Hungarian Academy of Sciences. HH is a Miguel Servet (CP14/00229) researcher funded by the Spanish Institute of Health Carlos III (ISCIII). This work has received funding from the Ministerio de Ciencia, Innovación y Universidades (SAF2017-89109-P; AEI/FEDER, UE).

## Author’s contributions

CHH, BS and JSR designed the research. CHH performed the analyses. BS, JT and JPP supervised by JSR and MPK and BAJ both supervised by DAL supported the development of the single cell simulation strategy. JT and JPP set up the cluster infrastructure for the simulation. JG supervised by OS processed the real scRNA-seq data. EM and HH provided the PBMC single cell data and supported the corresponding analysis. CHH, BS, JT, JG and JSR wrote the manuscript. JSR and BS supervised the project. All authors read, commented and approved the final manuscript.

## Competing interests

The authors declare that they have no competing interests

## Supplemental information Supplemental figures

**Fig. S1:**
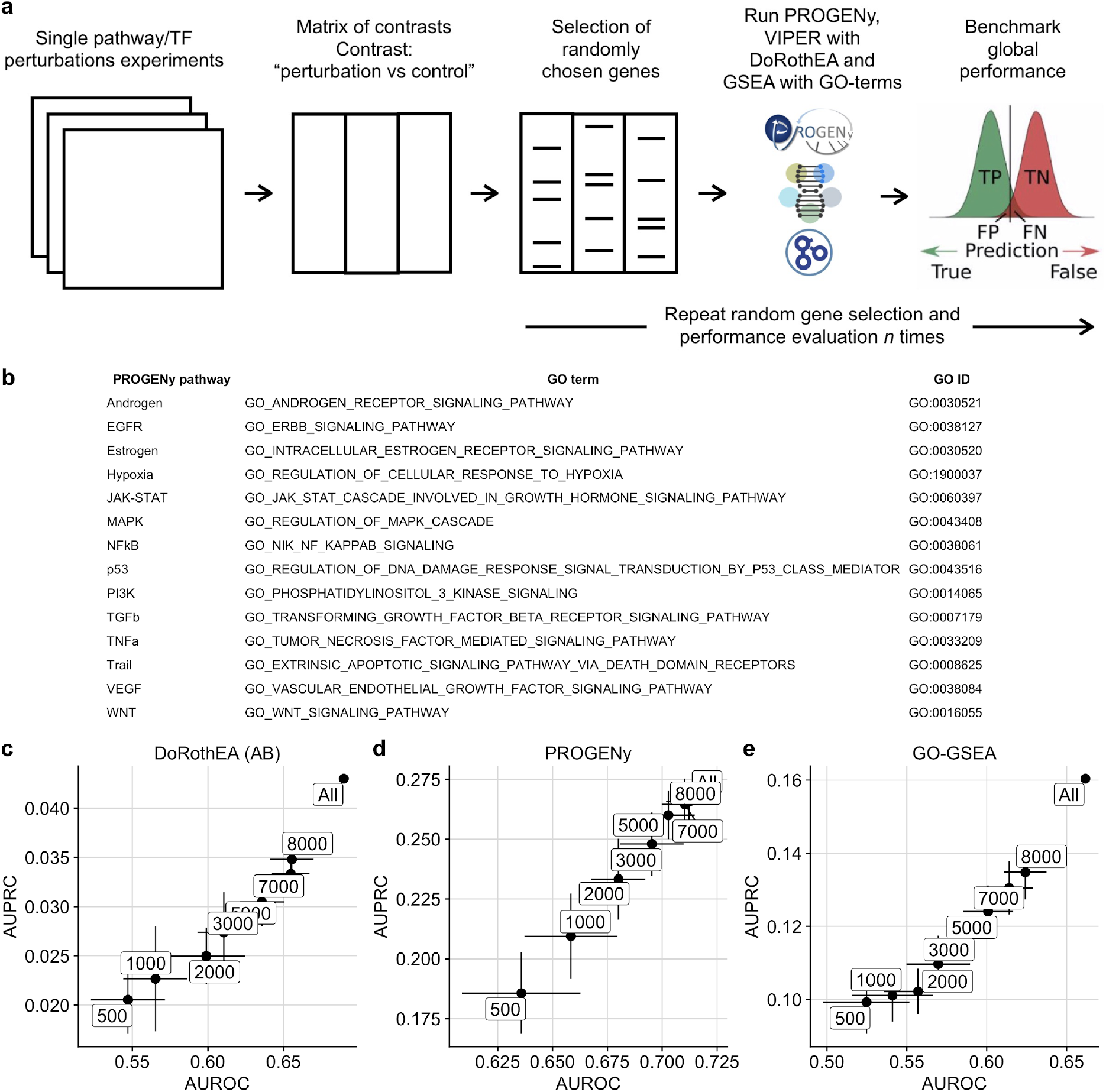
**a** Workflow to induce low gene coverage with subsequent benchmark. **b** Mapping table between PROGENy pathways and GO/GO IDs. **c,d,e** Scatterplot showing how well AUROC and AUPRC of **c** DoRothEA (AB) **d** PROGENy with 100 footprint genes per pathway and **e** GO-GSEA correspond to each other with respect to gene coverage.

**Fig. S2:**
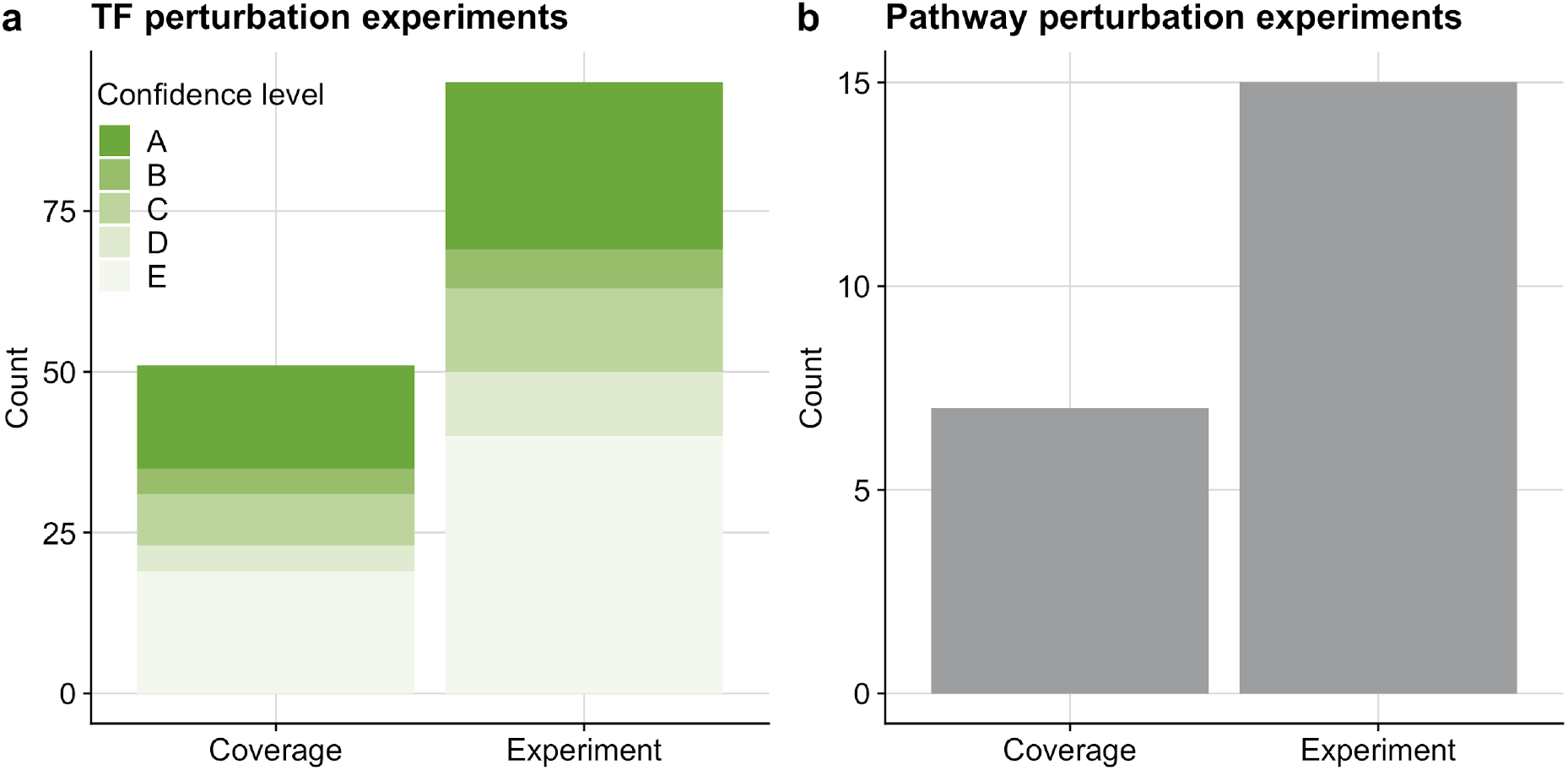
Overview of benchmark dataset for **a** TF activity and **b** Pathway activity inference tools.

**Fig. S3:**
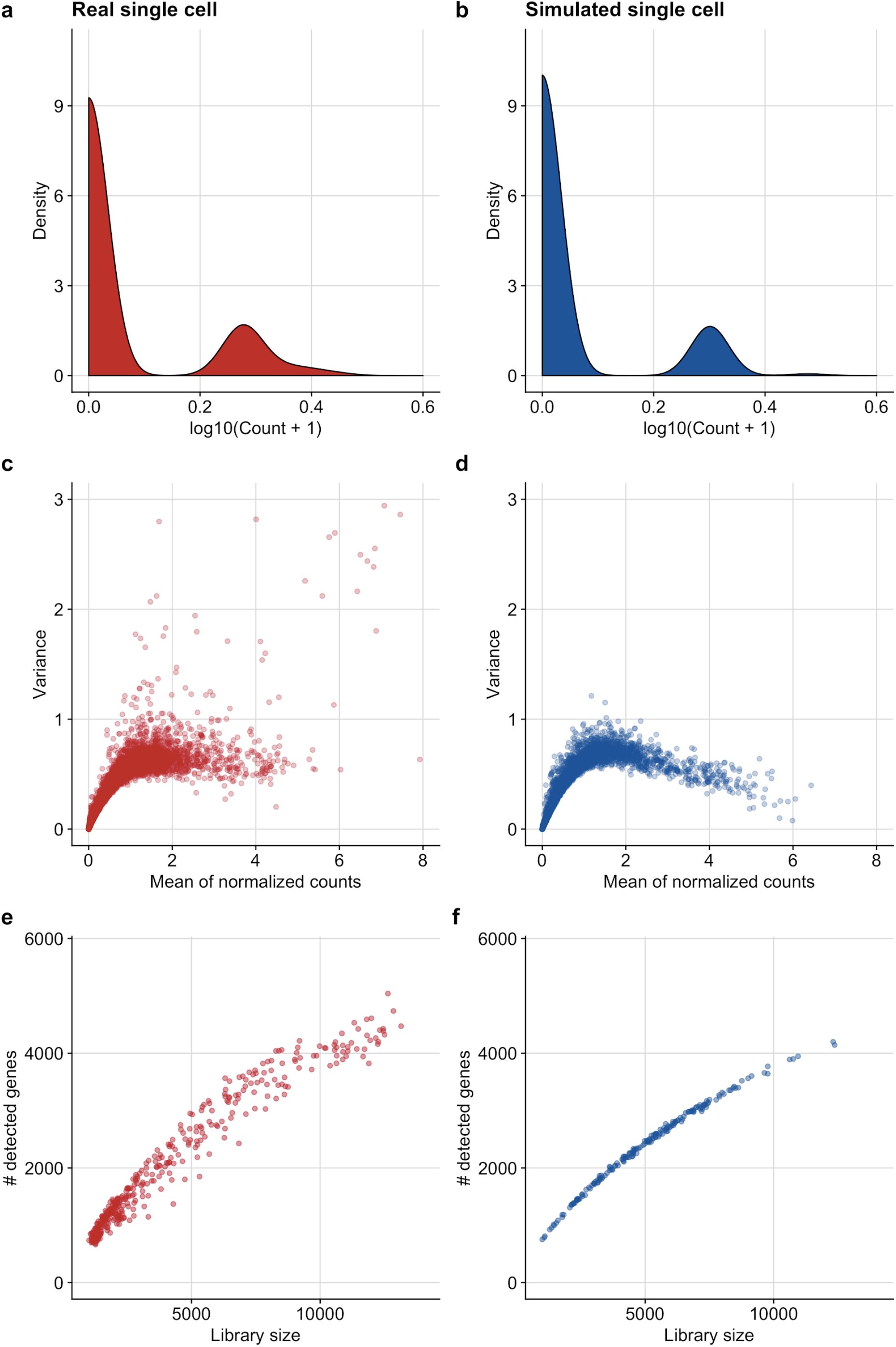
Comparison of single cell specific properties between real and simulated single cells. Count distribution of a representative gene for a **a** real and **b** simulated single cell. Relationship of mean to variance of a representative data set for a **c** real and **d** simulated single cell. Relationship of library size to number of detected genes for a **e** real and **f** simulated single cell.

**Fig. S4:**
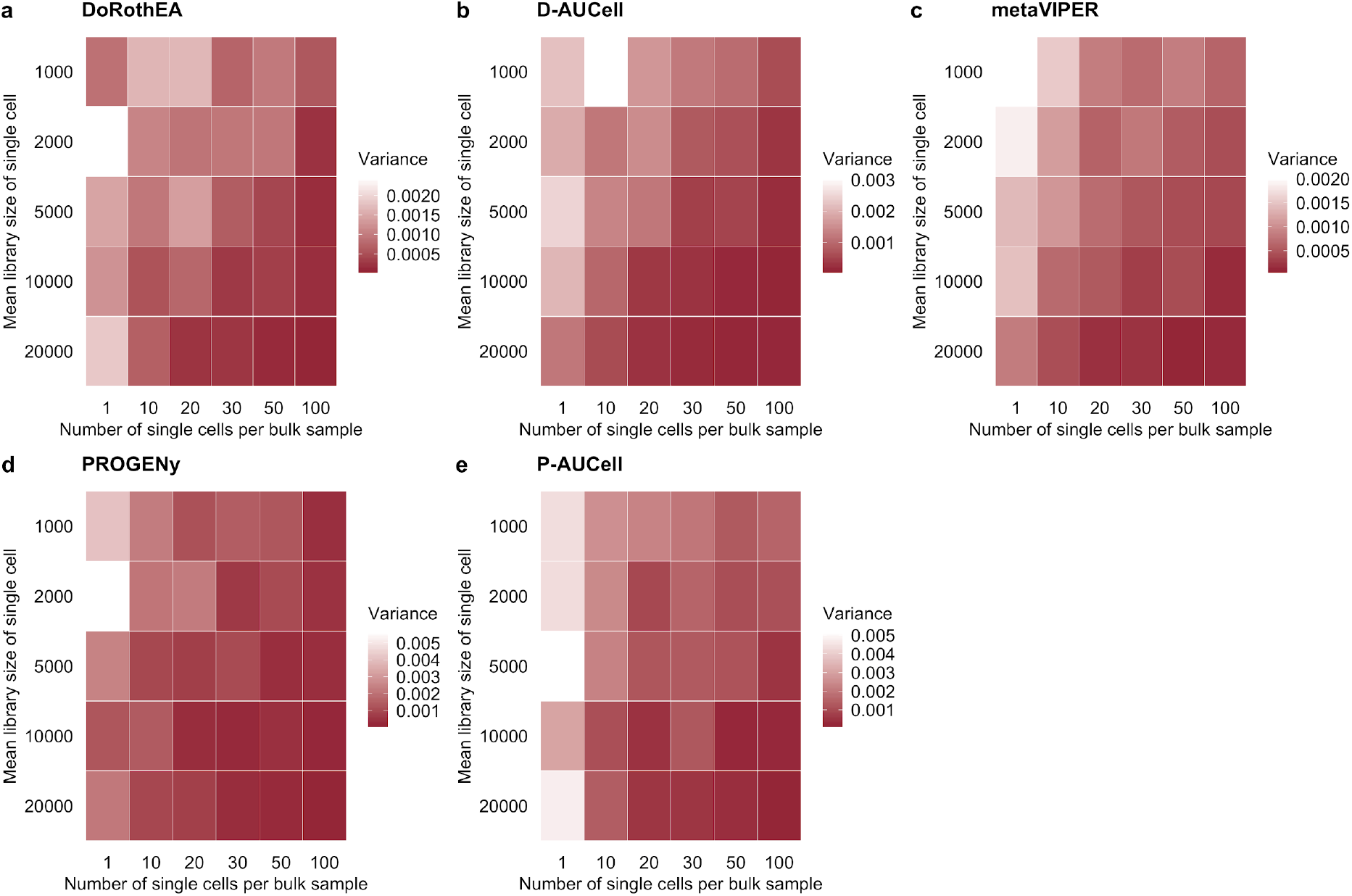
Variance of multiple performance evaluations (AUROC) of **a** DoRothEA, **b** D-AUCell, **c** metaVIPER, **d** PROGENy and **e** P-AUCell on single cells for different simulation parameter combinations.

**Fig. S5:**
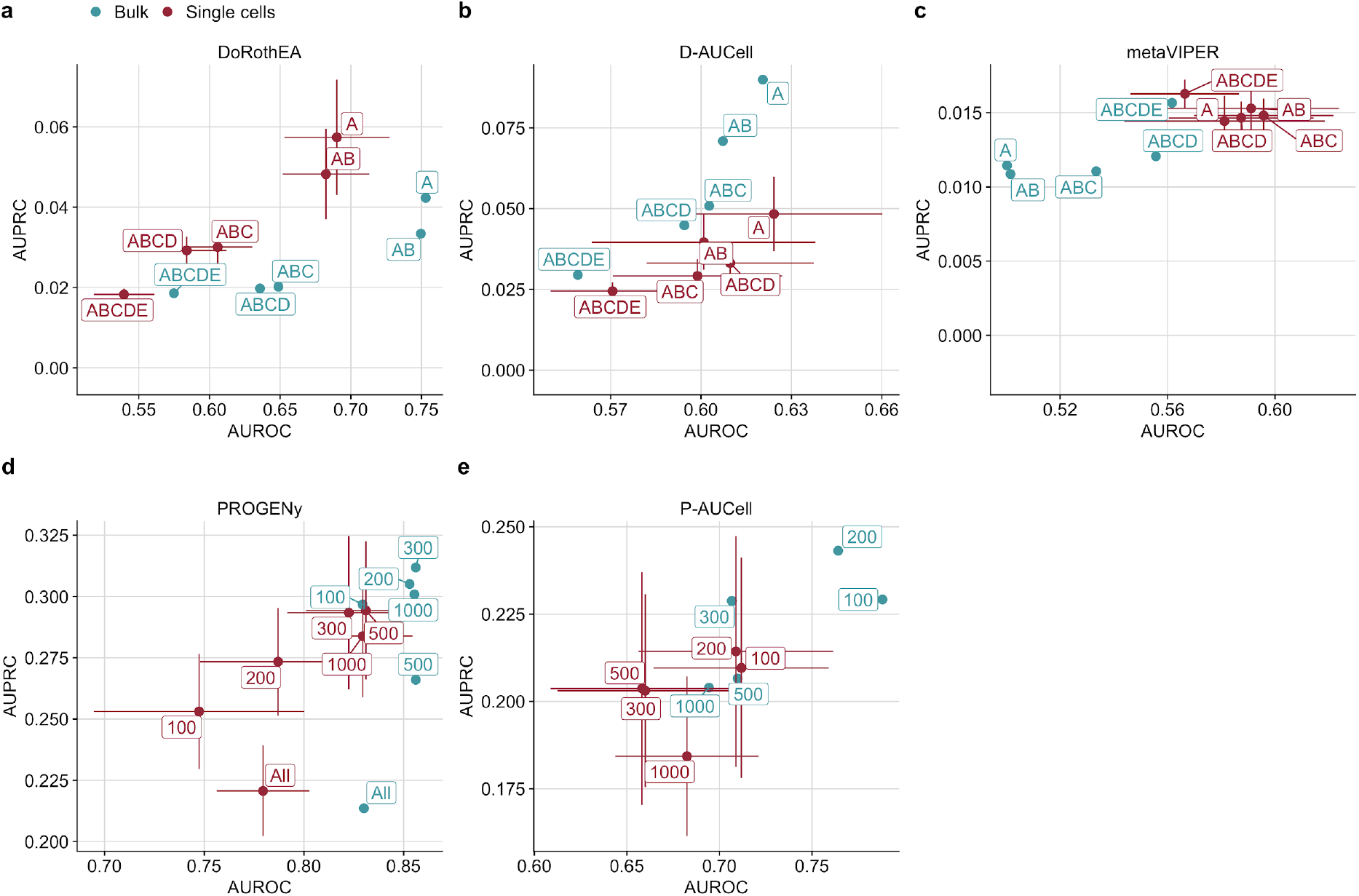
Scatterplot showing how well AUROC and AUPRC of **a** DoRothEA, **b** D-AUCell, **c** metaVIPER, **d** PROGENy and **e** P-AUCell performance on single cells and bulk correspond to each other with respect to different combinations of DoRothEA’s confidence levels or different number of footprint genes per pathway.

**Fig. S6:**
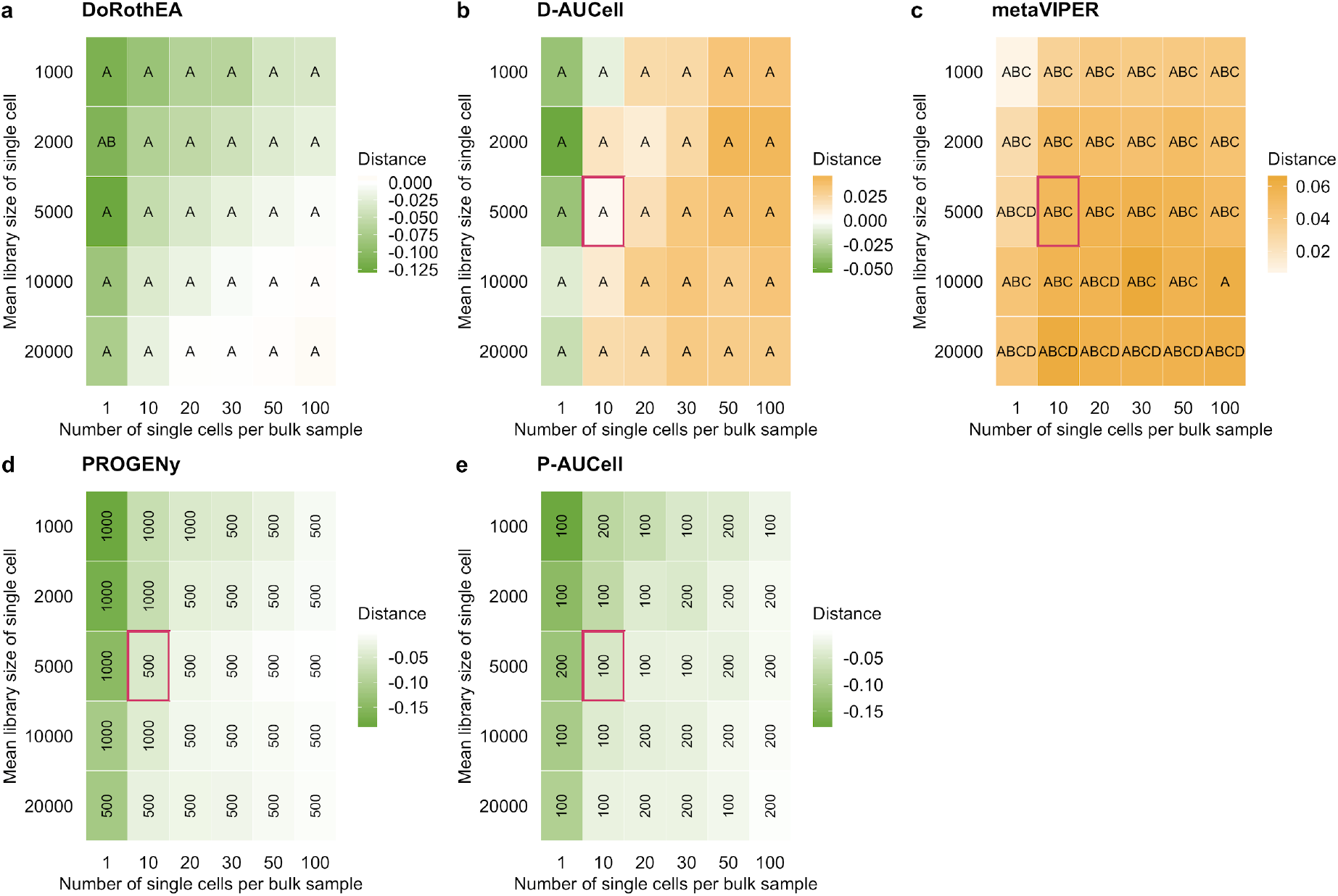
Distance heatmaps showing the performance difference of **a** DoRothEA, **b** D-AUCell, **c** metaVIPER, **d** PROGENy and **e** P-AUCell on single cells and corresponding bulk samples across all confidence level combinations or different number of footprint genes per pathway. The letters/numbers within the tiles indicates which confidence level combination/number of footprint genes per pathway performs the best. The tile marked in red corresponds to the parameter setting used for previous plots (Fig. 2 and Fig. S5)

**Fig. S7:**
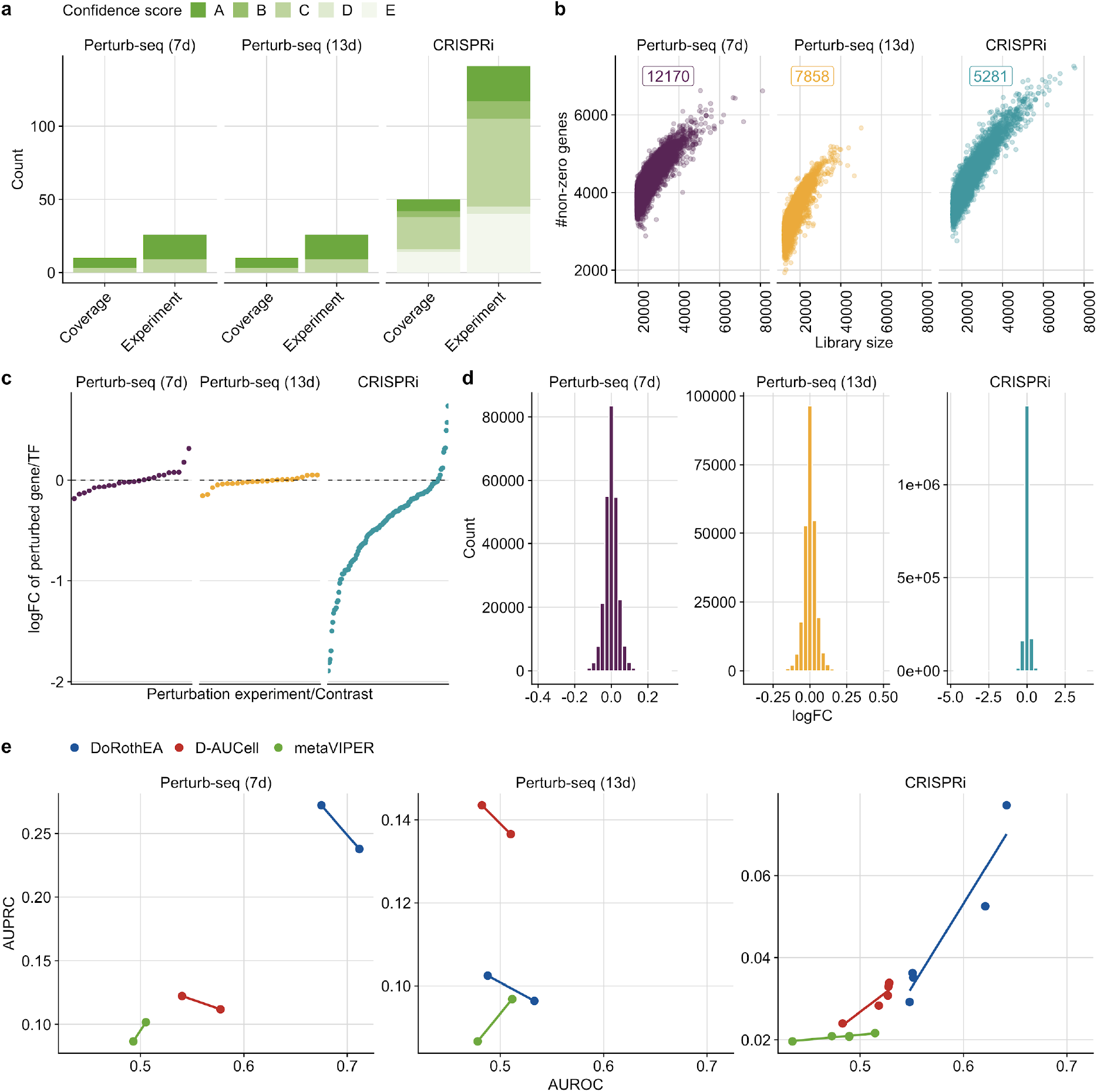
**a** Overview of benchmark dataset. **b** Relationship of library size to number of detected genes for all benchmark sub datasets. Number of corresponding cells are displayed as well. **c** logFC of perturbed target/TF for the corresponding perturbation experiment for all benchmark sub datasets. **d** Distribution of logFC for all genes and benchmark sub datasets. **e** Relationship between AUROC and AUPRC for all three methods with respect to different combinations of DoRothEA’s confidence levels for each sub benchmark dataset.

**Fig. S8.**
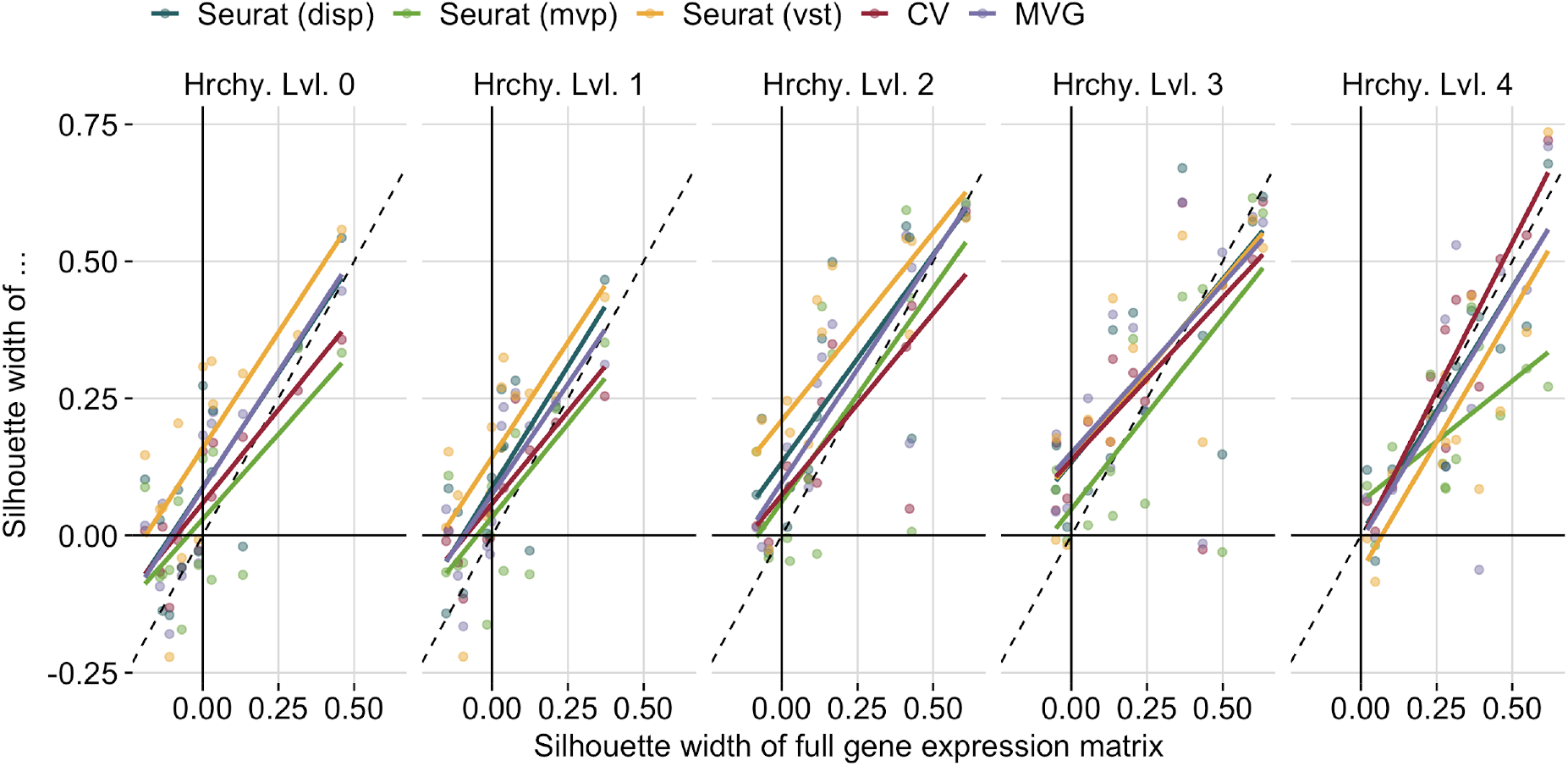
Identification of the best method to determine highly variable genes across single cells. We tested three different selection methods related to Seurat (disp = dispersion, mvp = mean.var.plot, vst). In addition we included CV (squared coefficient of variation - (sd/mean) **2) and MVG (most variable genes - genes with the highest variance).

**Fig. S9:**
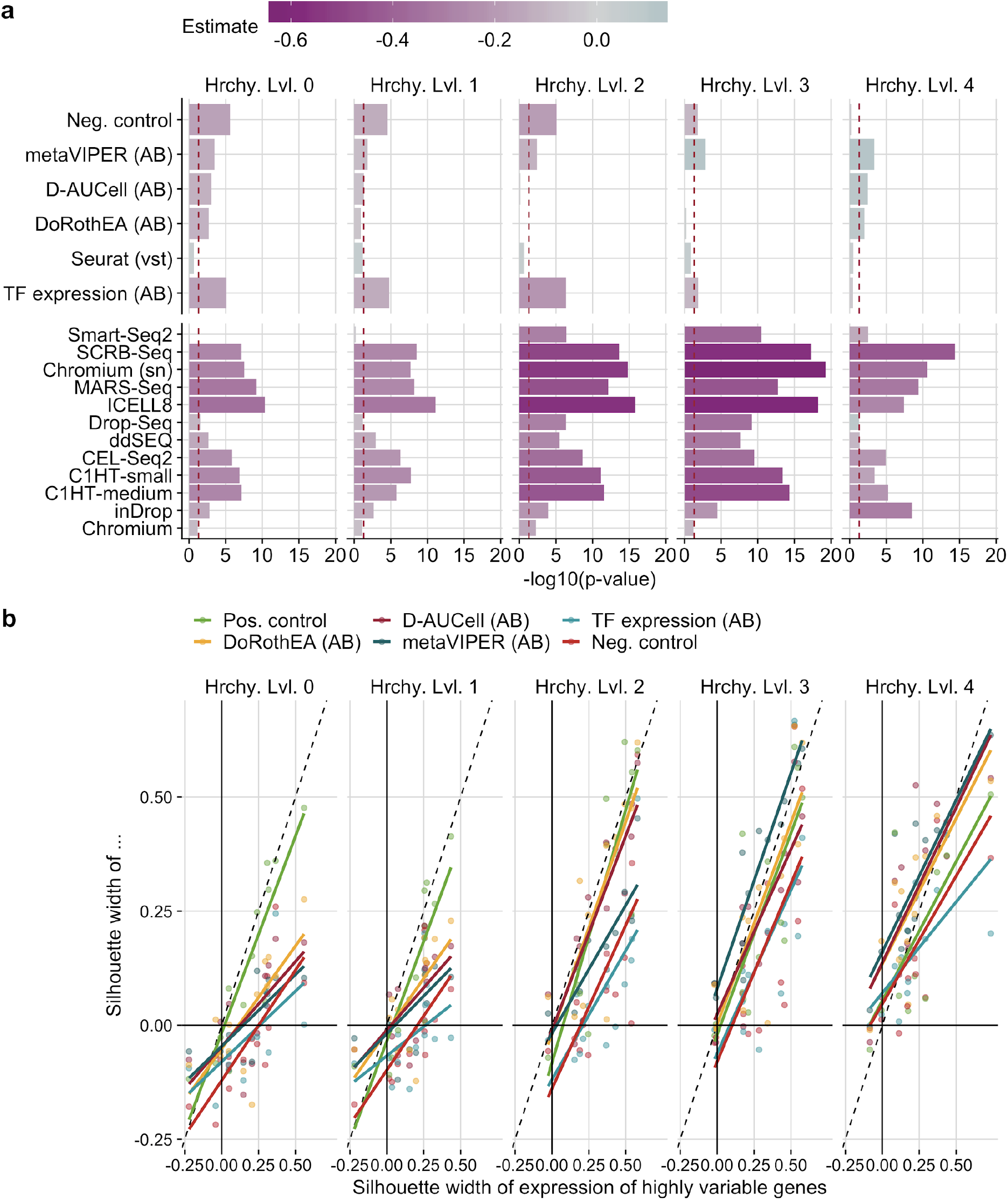
**a** Statistical analysis to evaluate two properties of the cluster purity analysis: i) whether different input matrices yield in better cluster purity than the positive control and ii) whether different scRNA-seq protocol yield in better cluster purity than Quartz-Seq2 for pathway activity inference tools. This analysis was performed independently for all hierarchy levels (Hrchy. Lvl.). Dashed line indicates a p-value of 0.05. **b** Comparison of cluster quality between HVGs and TF activity inference tool for all hierarchy levels (Hrchy. Lvl.).

**Fig. S10:**
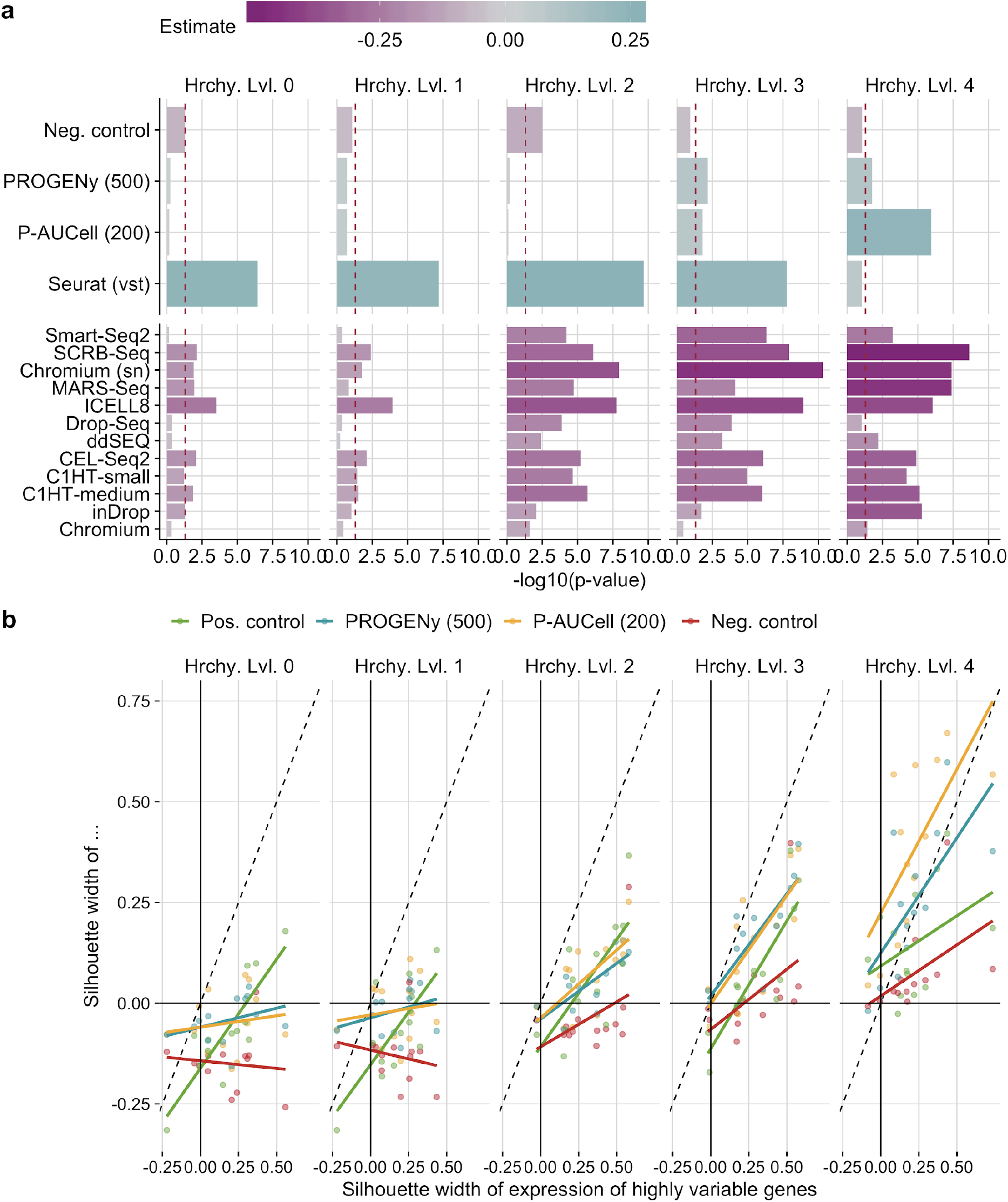
**a** Statistical analysis to evaluate two properties of the cluster purity analysis: i) whether different input matrices yield in better cluster purity than the positive control and ii) whether different scRNA-seq protocol yield in better cluster purity than Quartz-Seq2 for TF activity inference tools. This analysis was performed independently for all hierarchy levels (Hrchy. Lvl.). Dashed line indicates a p-value of 0.05. **b** Comparison of cluster quality between HVGs and pathway activity inference tools for all hierarchy levels (Hrchy. Lvl.).

**Fig. S11:**
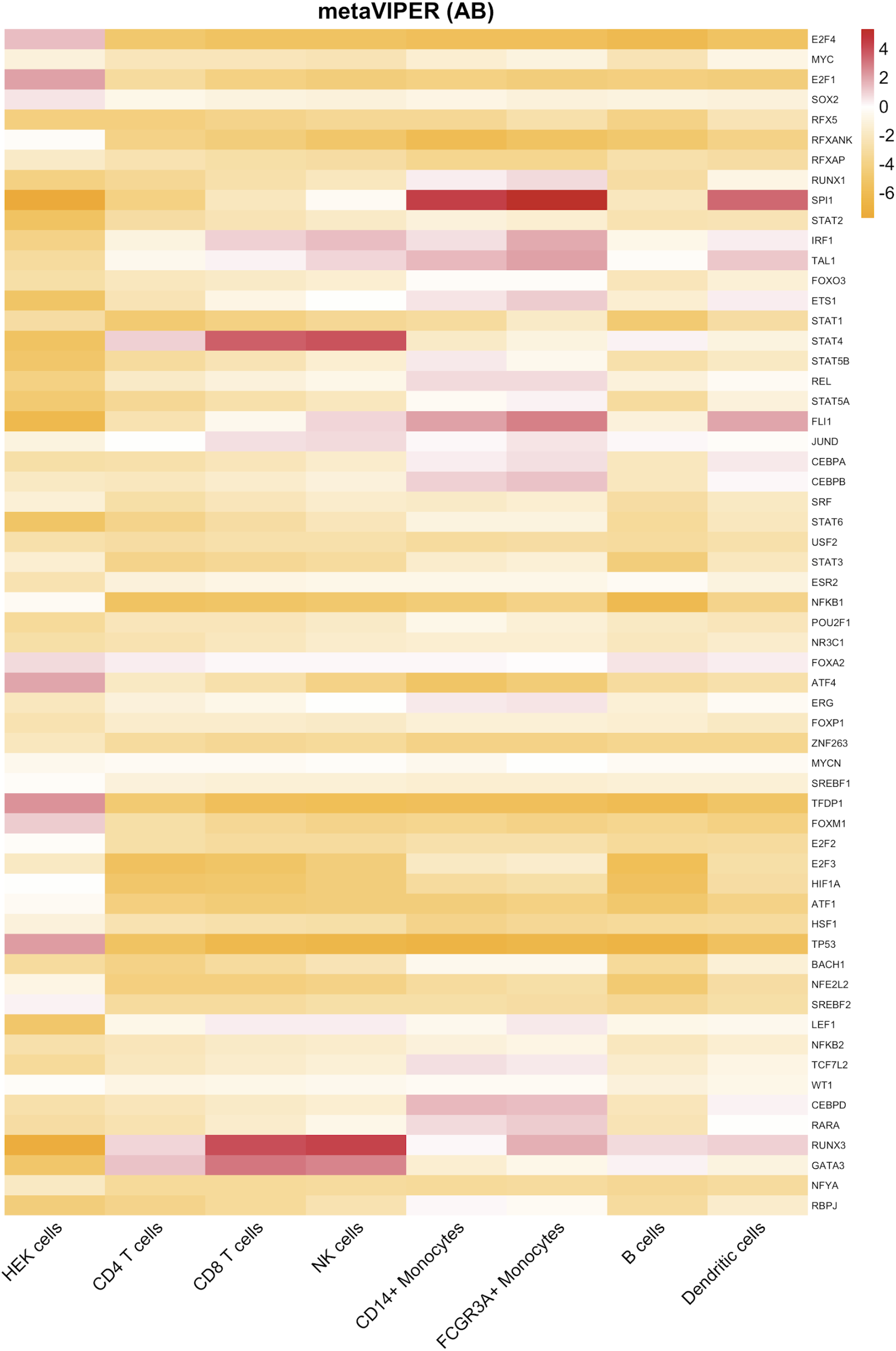
TF activities inferred from gene expression (Quartz-Seq2) using metaVIPER summarized for each cell type/cell line separately.

